# PCDH17 regulates lysosomal degradative capacity to promote autophagy attenuation during prolonged starvation

**DOI:** 10.64898/2026.09.03.749311

**Authors:** Bohan Chen, Shiou-Ling Lu, Takeshi Noda

## Abstract

Autophagy is induced by nutrient starvation to recycle intracellular constituents; however, its activity must subsequently be attenuated during prolonged nutrient deprivation. The mechanisms underlying this attenuation in mammalian cells remain incompletely understood. Here, using complementary HaloTag-based assays, we show that autophagic activity declines during prolonged starvation in HeLa cells. A genome-wide CRISPR/Cas9 knockout screen designed to identify cells that sustain autophagic activity under these conditions identified protocadherin 17 (PCDH17) as a regulator of autophagy attenuation.

PCDH17 depletion maintained autophagic activity during prolonged starvation without detectably altering mTORC1 signaling, ULK1 abundance, or the proximal machinery of autophagosome formation. Instead, PCDH17 depletion increased lysosomal abundance, acidification, and proteolytic activity, whereas PCDH17 overexpression produced reciprocal effects. We further identified a lysosome-associated PCDH17 subpopulation that is supplied predominantly through the biosynthetic ER–Golgi pathway. This pool undergoes proteolytic processing and lysosomal turnover, with starvation preferentially accelerating degradation of the C-terminal fragment while preserving a comparatively stable N-terminal fragment.

Together, these findings identify PCDH17 as an unexpected negative regulator of lysosomal function and demonstrate that modulation of lysosomal degradative capacity contributes to autophagy attenuation during prolonged starvation.

## Introduction

Cells maintain proteostasis and organelle quality through multiple intracellular degradation systems, most prominently the ubiquitin–proteasome system and lysosome-dependent pathways (Dikic, 2017). Macroautophagy and microautophagy are lysosome-dependent pathways that degrade cytoplasmic constituents through distinct membrane-trafficking mechanisms. During macroautophagy, cytoplasmic cargo is sequestered within double-membrane autophagosomes and subsequently delivered to lysosomes. In contrast, microautophagy involves the direct engulfment of cytoplasmic material through invagination of endosomal or lysosomal membranes (Yim and Mizushima, 2020; Schuck, 2020; Lu et al., 2025). Macroautophagy (hereafter referred to as autophagy) is an evolutionarily conserved process that is strongly induced by nutrient starvation (Nakatogawa, 2020).

Although intracellular degradation counterbalances biosynthesis, it is essential for cellular adaptation and survival. By recycling amino acids, lipids, and other metabolites, and by removing damaged proteins and organelles, degradative pathways preserve metabolic and proteostatic homeostasis. However, their activity must be tightly regulated: excessive or unrestrained autophagic degradation can deplete essential cellular constituents, disrupt cellular homeostasis, and compromise viability (Liu et al., 2023). Understanding how autophagic degradation is activated and subsequently attenuated is therefore central to understanding cellular adaptation to nutrient stress.

The duration and magnitude of autophagic flux are also likely to be pathophysiologically important. Autophagy is essential for tissue homeostasis and is implicated in a broad range of diseases, including cancer and neurodegenerative disorders (Yamamoto et al., 2023). During persistent metabolic stress, autophagy can initially support survival by recycling intracellular constituents; however, continued degradation in the absence of appropriate attenuation may erode essential cellular material and compromise viability. Conversely, premature attenuation can impair proteostasis and organelle quality control. Thus, mechanisms that appropriately limit autophagic activity after its adaptive phase may influence cell survival and disease progression during chronic stress.

Nutrient starvation induces autophagy primarily through inhibition of mTORC1, thereby releasing the autophagy-initiation machinery from nutrient-dependent suppression (Noda and Ohsumi, 1998). Under certain starvation conditions, amino acids generated by autophagic degradation can reactivate mTORC1 and thereby contribute to autophagy termination (Yu et al., 2010). In addition, prolonged starvation has been reported to reduce the abundance of ULK1, a key kinase required for autophagy initiation (Allavena et al., 2016; Liu et al., 2016). However, ULK1 expression can recover during prolonged starvation in other settings (Nazio et al., 2016), suggesting that the mechanisms governing autophagy attenuation are context-dependent and remain incompletely understood. In our previous study in *Saccharomyces cerevisiae*, we identified Tag1 as a critical regulator of autophagy attenuation during prolonged starvation (Kira et al., 2021). In yeast, autophagic activity declines approximately 9–12 h after starvation onset, demonstrating that this process is actively regulated rather than simply reflecting passive nutrient depletion. However, no mammalian ortholog of Tag1 has been identified. We therefore sought to identify mammalian factors that regulate autophagic activity during prolonged starvation, particularly those acting through mechanisms distinct from mTORC1 reactivation and ULK1 downregulation.

To address this question, we performed a genome-wide CRISPR/Cas9 knockout screen coupled with fluorescence-activated cell sorting (FACS) in HeLa cells and identified protocadherin 17 (PCDH17) as a candidate regulator. Cadherins comprise a large superfamily of calcium-dependent cell-surface adhesion proteins that mediate cell–cell recognition and tissue organization. Their extracellular regions contain tandem cadherin repeats that support adhesive interactions, whereas their cytoplasmic domains can couple adhesion complexes to intracellular signaling and the cytoskeleton (Takeichi, 1991; Gumbiner, 2005). Protocadherins constitute a major subgroup of the cadherin superfamily. Like classical cadherins, protocadherins are single-pass transmembrane proteins with extracellular cadherin repeats that mediate calcium-dependent, predominantly homophilic interactions. However, protocadherins are structurally and functionally distinct from classical cadherins. They generally possess a greater number of extracellular cadherin repeats and cytoplasmic domains that lack the conserved catenin-binding motifs found in classical cadherins, suggesting that they engage distinct intracellular signaling and trafficking mechanisms (Yagi, 2008; Redies et al., 2005).

PCDH17 is a member of the δ-protocadherin subfamily within the protocadherin family (Yagi, 2008; Redies et al., 2005). Like other protocadherins, PCDH17 contains multiple extracellular cadherin repeats, a single transmembrane domain, and a cytoplasmic region distinct from those of classical cadherins. Although protocadherins are best known for their roles in cell–cell adhesion and neural circuit development, their divergent cytoplasmic domains raise the possibility of functions beyond adhesion at the plasma membrane.

PCDH17 is expressed prominently in the nervous system and has been implicated in neuronal development (Hoshina et al., 2013; Hayashi et al., 2014; Asakawa and Kawakami, 2018). It has also been associated with tumorigenesis in several tissues, suggesting functions beyond the nervous system (Costa et al., 2011; Liu et al., 2019; Haruki et al., 2010; Wu et al., 2015; Yin et al., 2016; Liu et al., 2024). Although protocadherins are best known for mediating cell–cell adhesion at the plasma membrane, emerging evidence indicates that they can also participate in intracellular membrane trafficking. In particular, γ-protocadherins have been implicated in endolysosomal trafficking and LC3-associated membrane tubulation (Hanson et al., 2010; O’Leary et al., 2011; Shonubi et al., 2015).

Here, we identify PCDH17 as a negative regulator of sustained autophagic activity during prolonged starvation. We show that PCDH17 regulates lysosomal abundance, acidification, and proteolytic capacity, without detectably altering mTORC1 signaling or the proximal machinery of autophagosome formation. We further characterize a lysosome-associated PCDH17 subpopulation that is supplied through the biosynthetic ER–Golgi pathway and undergoes proteolytic processing and lysosomal turnover. These findings identify an unexpected intracellular function of PCDH17 and support a model in which modulation of lysosomal degradative capacity contributes to autophagy attenuation during prolonged starvation.

## Results

### Autophagy is attenuated during prolonged starvation in HeLa cells

We first investigated how autophagic flux changes during prolonged starvation. To monitor flux, we employed the HaloTag-based autophagy assay. In this assay, a cytosolic HaloTag fusion protein is delivered to lysosomes by autophagy, where most of the fusion protein is degraded. By contrast, the HaloTag7 moiety, when covalently bound to a HaloTag ligand, is stabilized against lysosomal proteolysis and accumulates as a free HaloTag fragment (Park and Marqusee, 2005; Yim et al., 2022). Thus, immunoblot-based quantification of the free HaloTag fragment relative to the total amount of free and full-length HaloTag species provides a cumulative measure of autophagic delivery and degradation within lysosomes (Yim et al., 2022).

HeLa cells stably expressing HaloTag7–mGFP were pulse-labeled with a HaloTag ligand in nutrient-rich medium for 20 min and subsequently starved for up to 36 h. Cells were harvested at the indicated time points and analyzed by immunoblotting (Fig. 1A). The ratio of the accumulated free HaloTag7 fragment to the total HaloTag7–mGFP-derived signal, defined as the sum of full-length HaloTag7–mGFP and the free HaloTag7 fragment, increased during the early phase of starvation but reached a plateau after approximately 9–12 h (Fig. 1B). Because this assay provides a cumulative measure of autophagic degradation, the plateau suggested that the rate of ongoing autophagic degradation declined during prolonged starvation.

**Figure 1.**
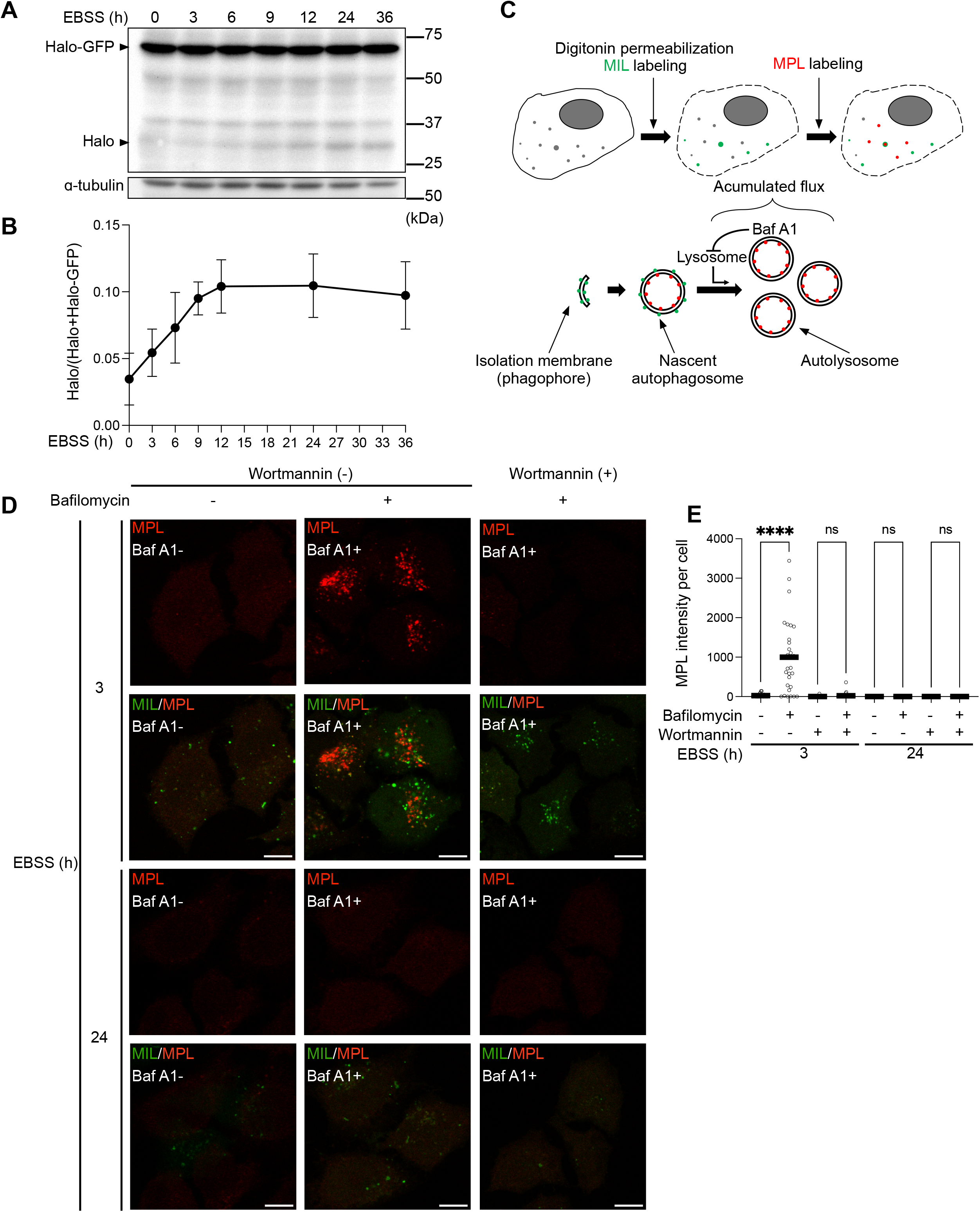
Autophagic activity declines during prolonged starvation in HeLa cells. (A) HeLa cells stably expressing HaloTag7-mGFP were pulse-labeled with 100 nM tetramethylrhodamine (TMR) HaloTag ligand in DMEM for 20 min at 37°C. After labeling, cells were washed and incubated in ligand-free EBSS for the indicated chase periods. Cell lysates were analyzed by immunoblotting with antibody against HaloTag and the indicated loading control. A representative immunoblot showing full-length HaloTag7-mGFP and the free HaloTag fragment is presented. (B) Quantification of HaloTag processing in (A). The proportion of free HaloTag was calculated as the intensity of the free HaloTag fragment divided by the combined intensities of free HaloTag and full-length HaloTag7-mGFP. Data are presented as the mean ± SD of three independent experiments. (C) Schematic representation of the Takahashi assay. HeLa cells expressing HaloTag7-LC3 were selectively permeabilized with digitonin and sequentially labeled with a membrane-impermeant Alexa Fluor 488 HaloTag ligand (MIL) and a membrane-permeant TMR HaloTag ligand (MPL). Where indicated, cells were treated with 200 nM bafilomycin A1 for 1 h before ligand labeling. (D, E) HeLa cells expressing HaloTag7-LC3 were starved in EBSS for 3 or 24 h. During the final 1 h of starvation, cells were treated with 200 nM bafilomycin A1 in the presence or absence of 200 nM wortmannin and subsequently subjected to the Takahashi assay. (D) Representative confocal images of MPL fluorescence. Scale bars, 10 μm. (E) Quantification of MPL fluorescence intensity per cell. Statistical significance was assessed by one-way ANOVA (\*\*\*\**p* < 0.0001; n > 30 cells per condition).

To directly evaluate autophagic flux over a defined interval, we next employed a HaloTag7–LC3-based assay, hereafter referred to as the Takahashi assay (Fig. 1C; Takahashi et al., 2018, 2019). LC3, a mammalian homolog of yeast Atg8, is synthesized as the soluble cytosolic form LC3-I and is subsequently lipidated with phosphatidylethanolamine to generate the membrane-associated form LC3-II during autophagosome formation (Kabeya et al., 2000; Ichimura et al., 2000). Following autophagosome closure, LC3-II on the cytosolic surface of the outer autophagosomal membrane is delipidated by ATG4 and recycled to the cytosol, whereas LC3-II associated with the inner membrane remains sequestered within the autophagosome and is subsequently delivered to autolysosomes for degradation (Kirisako et al., 2000; Hemelaar et al., 2003). The Takahashi assay exploits this membrane topology to distinguish cytosol-accessible HaloTag7–LC3 from HaloTag7–LC3 enclosed within autophagic compartments.

HeLa cells stably expressing HaloTag7–LC3 were starved to induce autophagy and then mildly permeabilized with digitonin, which selectively permeabilizes the plasma membrane while preserving intracellular organelle membranes (Fig. 1C). This procedure releases soluble cytosolic HaloTag7–LC3-I. The remaining cells were first incubated with a **<u>m</u>**embrane-**i**mpermeant HaloTag **l**igand conjugated to Alexa Fluor 488, hereafter designated **<u>MIL</u>**. Following plasma membrane permeabilization, MIL labels HaloTag7–LC3 that remains accessible from the cytosol, including LC3-II associated with phagophores/isolation membrane and the cytosolic surface of autophagosomes.

The cells were subsequently incubated with a **<u>m</u>**embrane-**<u>p</u>**ermeant HaloTag **l**igand conjugated to tetramethylrhodamine, hereafter designated **<u>MPL</u>**. Because cytosol-accessible HaloTag7–LC3 had already reacted with MIL, MPL preferentially labels the remaining HaloTag7–LC3 protected from MIL by intact organelle membranes, principally LC3-II enclosed within sealed autophagosomes and autolysosomes. Bafilomycin A1 inhibits lysosomal acidification and degradation (Yoshimori et al., 1991), causing membrane-enclosed HaloTag7–LC3 to accumulate during the treatment period. Accordingly, the bafilomycin A1-dependent increase in MPL fluorescence during the final 1 h of starvation provides a measure of autophagic flux during that defined interval (Fig. 1C).

After 3 h of starvation, treatment with bafilomycin A1 during the final 1 h markedly increased MPL fluorescence, and this increase was abolished by wortmannin, an inhibitor of autophagosome formation (Blommaart et al., 1997) (Fig. 1D and E). These results confirmed that the bafilomycin A1-dependent accumulation of MPL-labeled HaloTag7–LC3 reflects ongoing autophagic flux. By contrast, after 24 h of starvation, the increase in MPL fluorescence elicited by the same 1-h bafilomycin A1 treatment was significantly smaller than that observed after 3 h of starvation (Fig. 1D and E). Thus, two complementary HaloTag-based assays demonstrated that autophagic activity declines during prolonged starvation in HeLa cells.

### Genome-wide CRISPR screening identifies PCDH17 as a mediator of autophagy attenuation during prolonged starvation

We next adapted the Takahashi assay for flow-cytometric analysis. HeLa cells stably expressing HaloTag7–LC3 were starved for 3 h and treated with bafilomycin A1 during the final 1 h of starvation before sequential MIL/MPL labeling and flow-cytometric analysis. Wortmannin markedly reduced MPL fluorescence at comparable MIL fluorescence intensities, indicating that MPL fluorescence reflects ongoing autophagic flux (Fig. 2A). We next compared the MPL-to-MIL fluorescence ratio after 3 and 24 h of starvation. This ratio was markedly reduced after 24 h of starvation, consistent with attenuation of autophagic activity during prolonged starvation (Fig. 2B).

**Figure 2.**
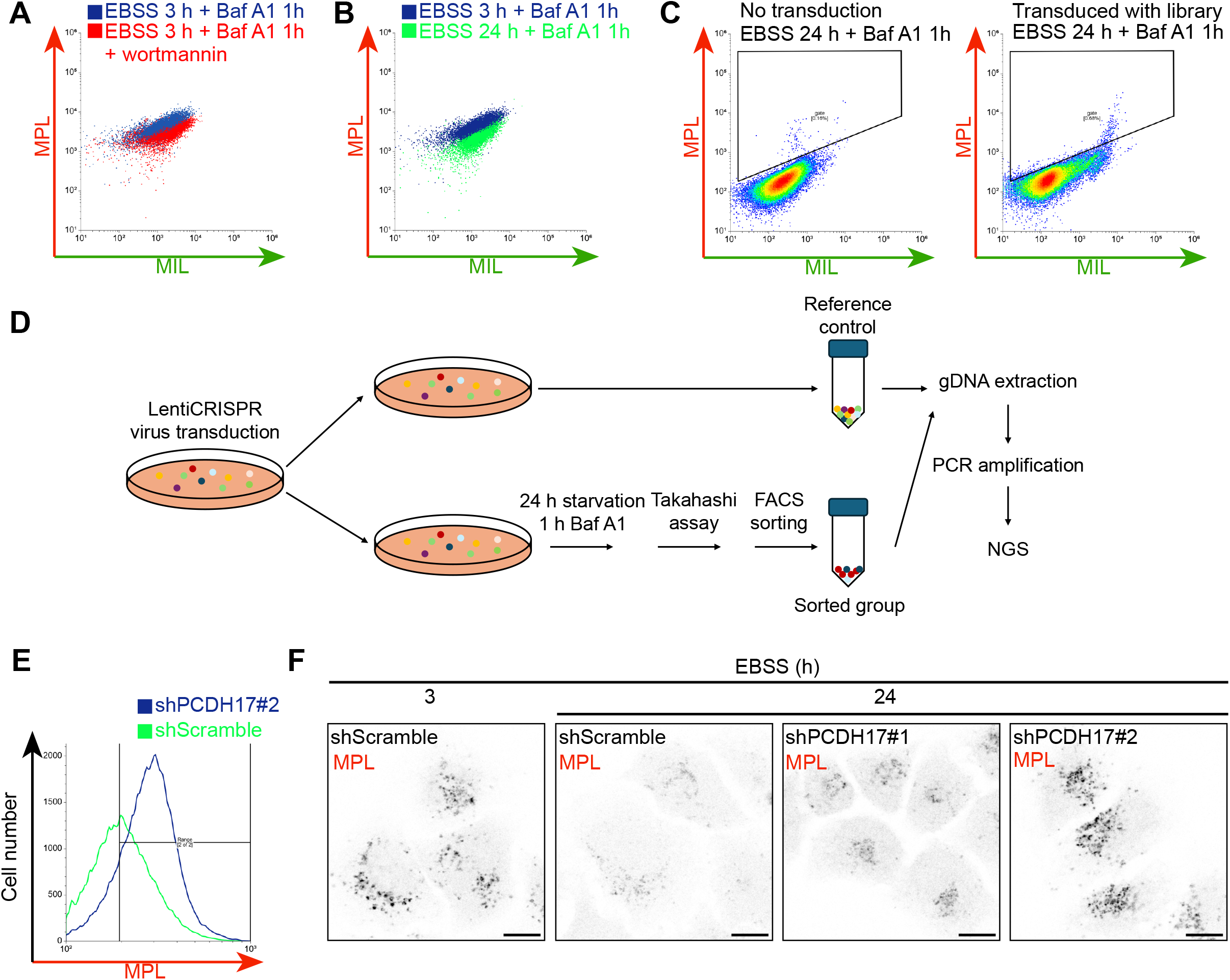
A genome-wide CRISPR screen identifies PCDH17 as a regulator of autophagy attenuation during prolonged starvation. (A, B) HeLa cells expressing HaloTag7-LC3 were starved in EBSS for 3 or 24 h and treated with 200 nM bafilomycin A1 during the final 1 h of starvation. Cells were then selectively permeabilized with digitonin and sequentially labeled with a membrane-impermeant Alexa Fluor 488 HaloTag ligand (MIL) and a membrane-permeant TMR HaloTag ligand (MPL) using the Takahashi assay. Representative flow cytometry profiles of MIL and MPL fluorescence following 3 h of starvation with or without wortmannin (A), and 3 or 24 h of starvation (B). (C) Representative MIL and MPL fluorescence profiles of non-library reference cells (left) and genome-wide CRISPR library-transduced cells (right) after 24 h of starvation, analyzed as described in (A). The sorting gate (black outline) was positioned outside the region occupied by the non-library reference population and then applied unchanged to the library-transduced population to isolate cells exhibiting increased MPL fluorescence. (D) Schematic overview of the genome-wide CRISPR screening workflow. (E) HeLa cells stably expressing HaloTag7-LC3 were transduced with lentiviruses encoding shPCDH17 #2 or a scrambled control shRNA. Following 24 h of starvation in EBSS, cells were subjected to the Takahashi assay and analyzed by flow cytometry. Representative histograms show cell counts plotted as a function of MPL fluorescence intensity. (F) HeLa cells stably expressing HaloTag7-LC3 were transduced with lentiviruses encoding shPCDH17 #1, shPCDH17 #2, or a scrambled control shRNA. Cells were starved in EBSS for 3 or 24 h and then subjected to the Takahashi assay. Representative confocal images of MPL fluorescence are shown. Scale bar, 10 μm.

We next performed a genome-wide CRISPR/Cas9 knockout screen in HeLa cells stably expressing HaloTag7–LC3. Cells transduced with the lentiviral sgRNA library were starved for 24 h, treated with bafilomycin A1 during the final 1 h of starvation, subjected to the Takahashi assay, and analyzed by flow cytometry (Fig. 2C, right). An autophagy-positive sorting gate was defined using the fluorescence profile of non-library-transduced control cells processed under the same starvation, bafilomycin A1 treatment, and ligand-labeling conditions. This gate selected cells with an elevated MPL-to-MIL fluorescence ratio, indicating sustained rather than attenuated autophagic activity after 24 h of starvation (Fig. 2C).

In parallel, an aliquot of the library-transduced cells was collected before starvation and was not exposed to bafilomycin A1, the Takahashi assay, or flow-cytometric sorting. This untreated library pool served as the reference population for subsequent sgRNA-enrichment analysis (Fig. 2C).

Genomic DNA was isolated from both the autophagy-positive sorted and reference populations, and sgRNA sequences were quantified to identify sgRNAs enriched in cells exhibiting sustained autophagic activity (Fig. 2D).

Across three independent screens, we selected 14 candidate genes that were enriched relative to the reference population in all three screens (*PCDH17, TULP1, SLC1A5, EEF1E1, CYSRT1, CYB561D2*) or were identified in more than three enriched sgRNA–screen events (*SESN1, HHLA3, FAM185A, LRP1B, VPS4B, ATG4D, FAM69B, PPAPDC1A*) across the combined datasets. In addition, four genes that showed robust enrichment in two of the three screens (*SENP1, ULK2, KCNJ5, MYRIP*) were included on the basis of their functional annotations, yielding a total of 18 candidates.

Each candidate was individually depleted in HeLa cells using lentivirally delivered shRNAs; for most genes, two independent shRNA clones were used (Supplemental table 2). After 24 h of starvation, knockdown cells were evaluated using the Takahashi assay coupled with flow cytometry and fluorescence microscopy, as well as the HaloTag-based autophagy assay (Fig. 2E and Fig. S1A–C). Among the candidates tested, PCDH17 depletion most consistently produced a phenotype indicative of sustained autophagic activity during prolonged starvation.

In polyclonal PCDH17-knockdown populations, in which PCDH17 protein abundance was reduced by approximately 50%, the number of MPL-positive HaloTag7–LC3 puncta was increased relative to scrambled-control cells after 24 h of starvation (Fig. 3A and B). To obtain homogeneous cell populations with sustained PCDH17 depletion, we subsequently isolated monoclonal knockdown cell lines by limiting dilution; these clones exhibited an approximately 90% reduction in PCDH17 expression (Fig. 3C). Consistent with the increased abundance of HaloTag7–LC3 puncta, monoclonal PCDH17-knockdown cells showed significantly enhanced degradation of HaloTag7–mGFP after 24 h of starvation (Fig. 3D and E). Together, these results support the conclusion that PCDH17 contributes to the attenuation of autophagic activity during prolonged starvation.

**Figure 3.**
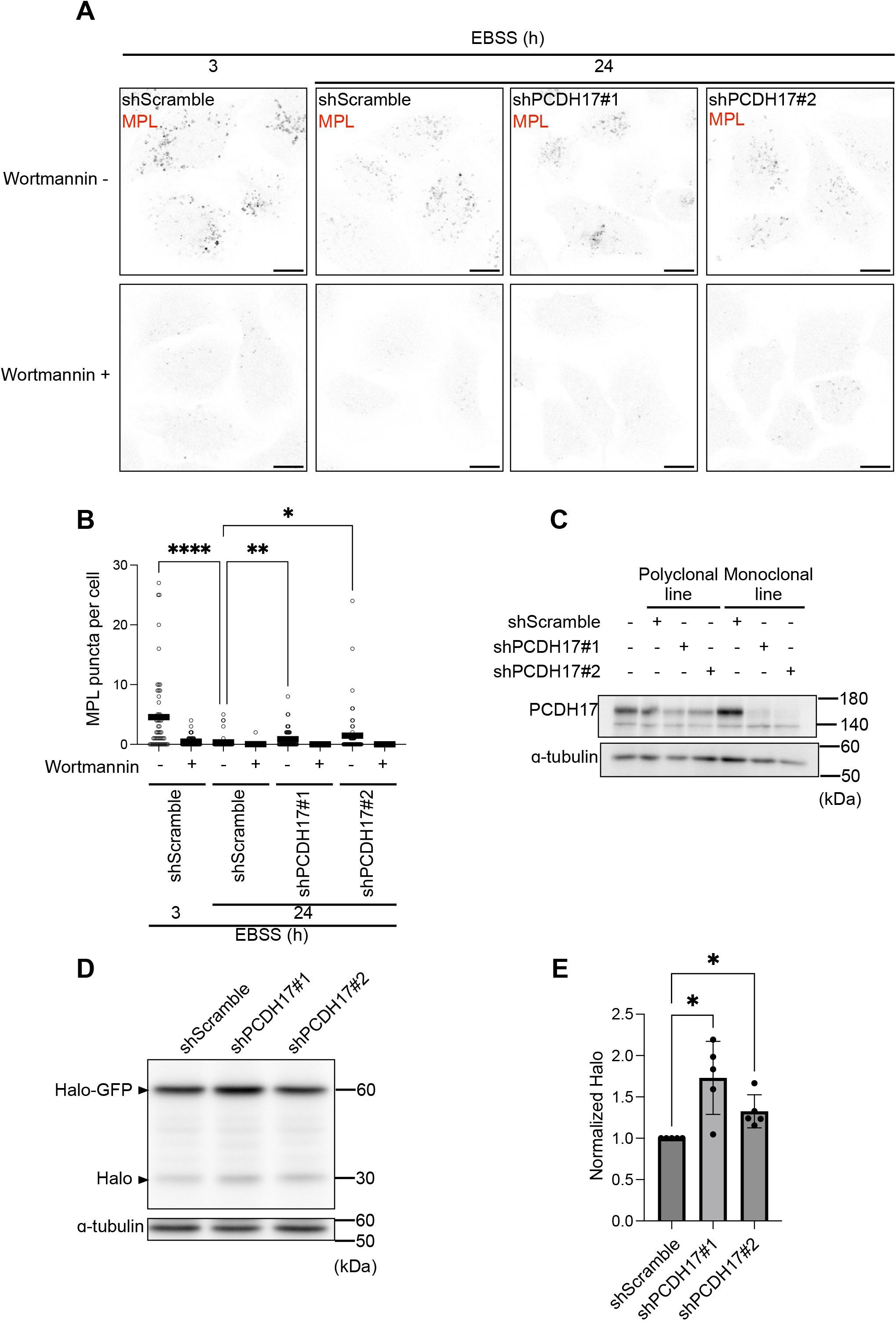
PCDH17 knockdown increases MPL-labeled HaloTag7-LC3 puncta and promotes HaloTag7-mGFP degradation. (A) HeLa cells stably expressing HaloTag7-LC3 were transduced with lentiviruses encoding shPCDH17 #1, shPCDH17 #2, or a scrambled control shRNA. Cells were starved in EBSS for 3 or 24 h, subjected to the Takahashi assay, and imaged by confocal microscopy. Representative images of MPL-labeled HaloTag7-LC3 puncta are shown. Scale bars, 10 μm. (B) Quantification of the number of MPL-labeled HaloTag7-LC3 puncta per cell in (A) (n > 30 cells per condition). Statistical significance was assessed by Kruskal-Wallis test. (C) Immunoblot analysis of PCDH17 expression in the indicated polyclonal and monoclonal PCDH17-knockdown cell lines. (D) Control and PCDH17-knockdown cells stably expressing HaloTag7-mGFP were pulse-labeled with 100 nM TMR HaloTag ligand for 20 min immediately before starvation in EBSS for the indicated durations. Cell lysates were analyzed by immunoblotting with the indicated antibodies. A representative immunoblot is shown. (E) Quantification of full-length HaloTag7-mGFP in (D), normalized to α-tubulin (n = 5 independent experiments). Data are presented as the mean ± SD. Statistical significance was assessed by one-way ANOVA. (\**p* < 0.05; ** *p* < 0.01; **** *p* < 0.0001).

### mTORC1 signaling and proximal autophagy machinery are largely unaffected by PCDH17 perturbation

Nutrient starvation induces autophagy primarily through inhibition of mTORC1, a central nutrient-responsive regulator of cellular metabolism and autophagy (Noda and Ohsumi, 1998). Under certain starvation conditions, autophagy-dependent amino acid replenishment can reactivate mTORC1, thereby contributing to autophagy termination (Yu et al., 2010). We therefore considered whether PCDH17 attenuates autophagy by promoting mTORC1 reactivation.

In our experimental conditions, however, phosphorylation of S6K, a canonical readout of mTORC1 activity, remained suppressed throughout prolonged starvation and did not recover even after 24 h in control cells (Fig. 4A). mTORC1 also phosphorylates ULK1 under nutrient-rich conditions, whereas starvation-induced mTORC1 inactivation promotes ULK1 dephosphorylation and autophagy initiation (Jung et al., 2009; Ganley et al., 2009; Hosokawa et al., 2009). Consistent with the S6K results, ULK1 rephosphorylation was not detected after 24 h of starvation in control cells (Fig. 4A). Thus, mTORC1 was not detectably reactivated during prolonged starvation under our experimental conditions.

**Figure 4.**
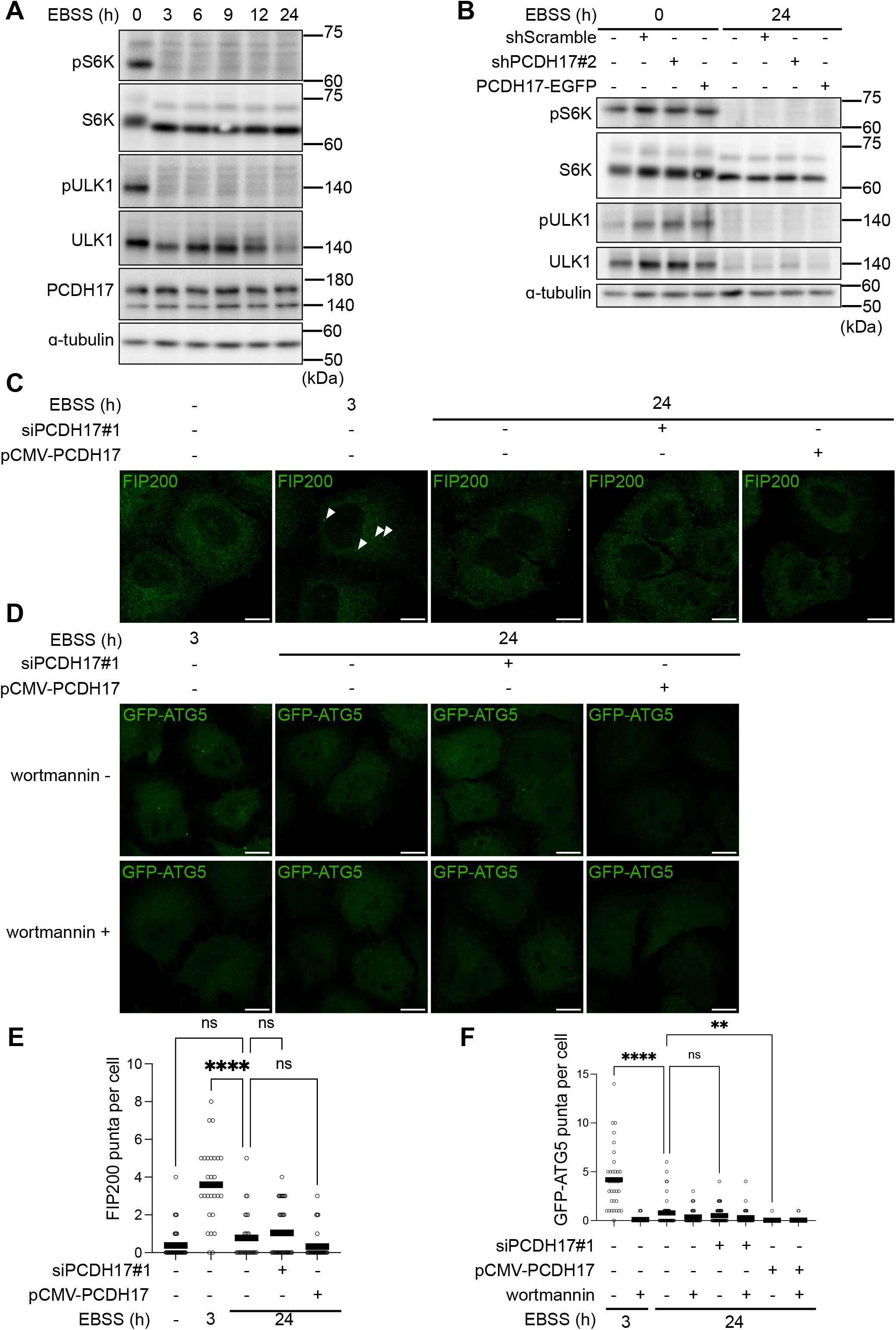
mTORC1 signaling and the upstream autophagy machinery are largely unaffected by PCDH17 depletion or overexpression. (A) HeLa cells were starved in EBSS for 0, 3, 6, 9, 12, or 24 h, and cell lysates were analyzed by immunoblotting with the indicated antibodies. (B) Control cells and cells expressing scrambled shRNA, shPCDH17 #2, or PCDH17–EGFP were maintained in complete medium or starved in EBSS for 24 h. Cell lysates were analyzed by immunoblotting with the indicated antibodies. (C– F) HeLa cells were transfected with siControl, siPCDH17 #1, empty pCMV vector, or pCMV-PCDH17 plasmid for 24 h and were starved in EBSS for 0, 3, 12 h and analyzed by confocal microscopy. (C) Cells were fixed and subjected to immunofluorescence staining with an anti-FIP200 antibody. (D) HeLa cells stably expressing GFP–ATG5 were imaged. Scale bars, 10 μm. (E, F) Quantification of FIP200- and GFP–ATG5-positive puncta per cell, respectively (n > 30 cells per condition). Data are presented as the mean ± SD. Statistical significance was assessed by Kruskal-Wallis test. (\*\**p* < 0.01; **** *p* < 0.0001).

Moreover, neither PCDH17 depletion nor PCDH17 overexpression affected the phosphorylation status of S6K or ULK1 (Fig. 4B and Fig. S2B), indicating that PCDH17-mediated autophagy attenuation is unlikely to result from altered mTORC1 signaling. Although total ULK1 abundance decreased markedly during prolonged starvation, consistent with previous reports linking ULK1 downregulation to autophagy termination (Liu et al., 2016; Allavena et al., 2016), PCDH17 knockdown did not alter this reduction (Fig. S2A). These findings suggest that PCDH17 regulates autophagy attenuation through a mechanism distinct from mTORC1 reactivation or ULK1 downregulation.

We next examined whether PCDH17 affects the proximal machinery for autophagosome formation. FIP200, together with ULK1 and ATG13, forms an autophagy-initiation complex that accumulates at sites of autophagosome biogenesis; accordingly, FIP200 puncta serve as an indicator of autophagosome initiation (Hara et al., 2008). After 24 h of starvation, the number of FIP200 puncta did not differ significantly between PCDH17-knockdown and control cells (Fig. 4C and E). Similarly, the number of ATG5-positive puncta, which reflect recruitment of the ATG12–ATG5–ATG16L1 complex to sites of LC3 lipidation and autophagosome biogenesis (Mizushima et al., 2001; Fujita et al., 2008), was unchanged in siPCDH17-treated cells relative to siControl-treated cells after 24 h of starvation (Fig. 4D and F; Fig. S2C). Collectively, these results do not support a major role for PCDH17 in regulating mTORC1 signaling or the upstream machinery of autophagosome formation during prolonged starvation.

### A lysosome-associated subpopulation of PCDH17 is dynamically regulated by nutrient availability

PCDH17 belongs to the protocadherin family, whose best-established function is to mediate cell– cell adhesion at the plasma membrane (Redies et al., 2005). We therefore examined the intracellular localization of PCDH17 under our experimental conditions (Fig. 5 and Fig. S3–S4). In addition to its expected plasma membrane localization, C-terminally GFP-tagged PCDH17 (PCDH17–GFP) exhibited a punctate cytoplasmic distribution under nutrient-rich conditions (Fig. 5A). These cytoplasmic puncta predominantly colocalized with the lysosomal marker LAMP1 in cells cultured in complete medium, indicating the presence of a lysosome-associated PCDH17 subpopulation in addition to the plasma membrane pool (Fig. 5B). Time-lapse imaging further showed that PCDH17–GFP puncta moved in concert with mRFP–LAMP1-positive structures (Video S1), supporting their association with the lysosomal compartment.

**Figure 5.**
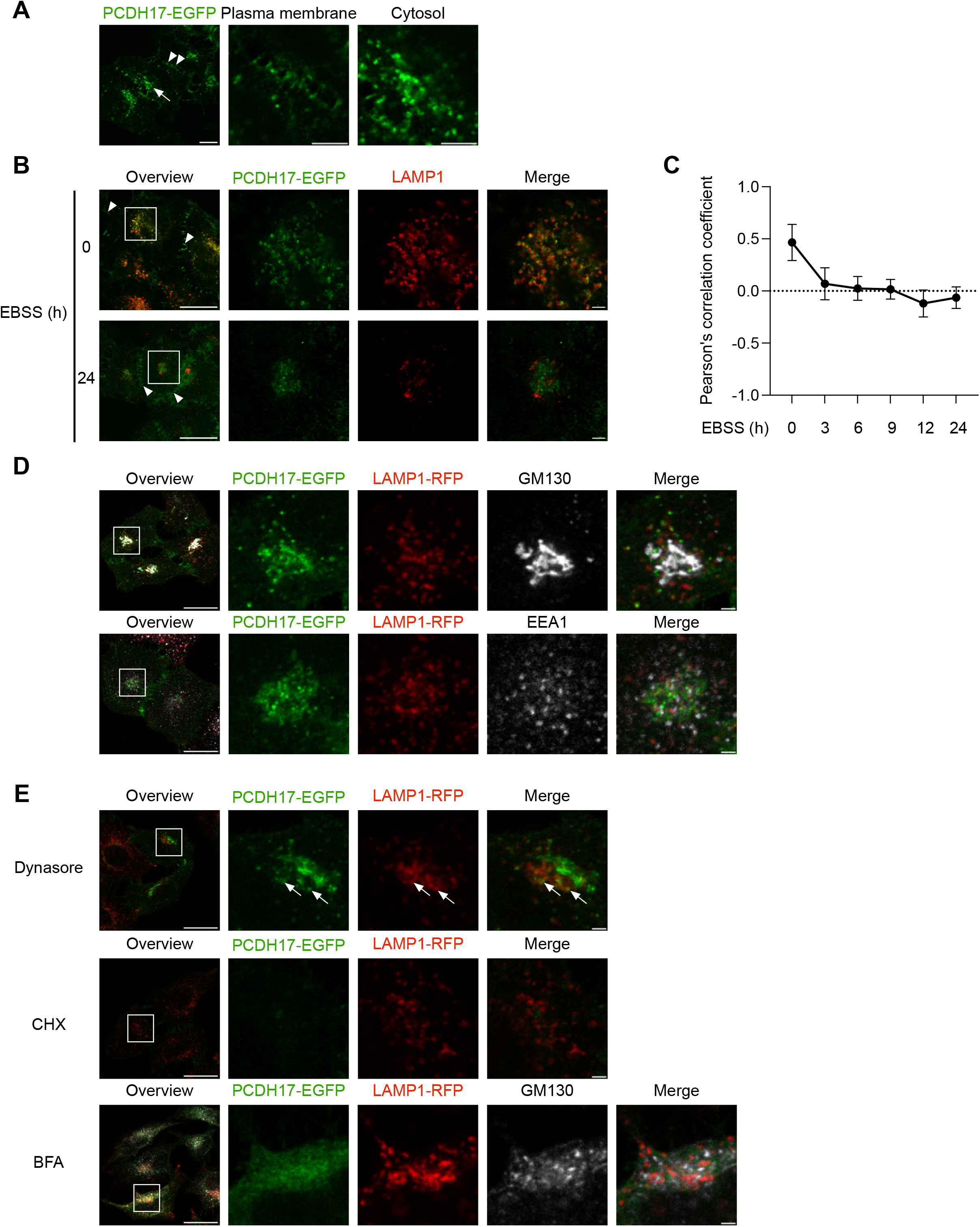
Dynamic localization of PCDH17 suggests its involvement in intracellular membrane trafficking. (A) HeLa cells expressing PCDH17–EGFP were cultured in complete medium and imaged by confocal microscopy. The middle and right panels show enlarged views of the indicated regions. White arrowheads indicate PCDH17–EGFP at the plasma membrane, whereas white arrows indicate cytoplasmic PCDH17–EGFP puncta. Scale bars, 10 μm in the overview and 5 μm in the enlarged images. (B) HeLa cells expressing PCDH17–EGFP were maintained in complete medium or starved in EBSS for 24 h, fixed, and immunostained for LAMP1. Boxed regions in the overview images are enlarged in the adjacent panels. White arrowheads indicate PCDH17–EGFP at the plasma membrane. The enlarged images were deconvolved using Huygens software. Scale bars, 20 μm in the overview images and 2 μm in the enlarged images. (C) Quantification of colocalization between PCDH17–EGFP and LAMP1 signals in (B). Pearson’s correlation coefficients were calculated using the Coloc 2 plugin in ImageJ with Costes automatic thresholding (n > 20 cells per condition). (D) HeLa cells expressing PCDH17–EGFP and LAMP1–RFP were starved in EBSS for 3 h and then returned to complete medium for 3 h. Cells were fixed, and stained with GM130 or EEA1 antibodies, and imaged by confocal microscopy. Boxed regions in the overview images are enlarged in the adjacent panels. Scale bars, 20 μm in the overview images and 2 μm in the enlarged images. (E) Following 3 h of starvation in EBSS, cells expressing PCDH17–EGFP and LAMP1–RFP were returned to complete medium for 3 h in the presence of either 80 μM dynasore, 5 μg/mL brefeldin A, or 50 μg/mL cycloheximide. Cells were fixed and imaged by confocal microscopy. Boxed regions in the overview images are enlarged in the adjacent panels. White arrows indicate lysosome-colocalized l PCDH17–EGFP puncta. Scale bars, 20 μm in the overview images and 2 μm in the enlarged images.

Following 3 h of starvation, however, PCDH17 was largely absent from LAMP1-positive lysosome, and its colocalization with LAMP1 was barely detectable after 24 h of starvation (Fig. 5B and C). By contrast, the plasma membrane abundance of PCDH17 remained unchanged (Fig. 5B; arrowheads). To determine whether the loss of lysosome-associated PCDH17 was reversible, cells starved for 3 h were returned to nutrient-rich DMEM. After 3 h of nutrient replenishment, most cytoplasmic PCDH17 puncta again colocalized with LAMP1, indicating that lysosomal localization of PCDH17 is reversible (Fig. 5D).

These observations raised two possibilities for the origin of lysosome-associated PCDH17: endocytosis of the plasma membrane pool or biosynthetic delivery of newly synthesized PCDH17 through the ER–Golgi pathway. To distinguish between these possibilities, we examined PCDH17 localization at an intermediate stage of nutrient replenishment. First lysosomal PCDH17 disappeared under starvation for 3 h, and after cells were returned to nutrient-rich DMEM for further 3 h, the cytoplasmic PCDH17 puncta reappeared, but showed little colocalization with LAMP1, suggesting that they represent intermediate en route to lysosomes (Fig. 5D). Instead, the newly formed PCDH17 puncta colocalized extensively with the Golgi marker GM130, but not with the early endosome marker EEA1 (Fig. 5D).

We next treated cells with dynasore, an inhibitor of dynamin-dependent endocytosis, during nutrient replenishment following 3 h of starvation (Macia et al., 2006). Prominent PCDH17 puncta still emerged after 3 h of replenishment despite dynasore treatment (Fig. 5E; arrows). In contrast, brefeldin A, which disrupts ER-to-Golgi trafficking (Misumi et al., 1986), markedly suppressed the reappearance of PCDH17 puncta (Fig. 5E). Under brefeldin A treatment, PCDH17 instead exhibited a reticular distribution and colocalized with dispersed GM130-positive structures, consistent with disruption of Golgi organization and retention of PCDH17 within the early secretory pathway. Inhibition of de novo protein synthesis with cycloheximide similarly abolished the reappearance of PCDH17 puncta (Fig. 5E). Together, these results indicate that lysosome-associated PCDH17 is derived from newly synthesized protein delivered through the ER–Golgi pathway, rather than through endocytosis of the plasma membrane pool.

### PCDH17 undergoes lysosomal degradation, particularly during starvation

As described above, lysosome-associated PCDH17–GFP fluorescence progressively disappeared during starvation. This loss was prevented by bafilomycin A1, an inhibitor of vacuolar H⁺-ATPase (V-ATPase) (Fig. 6A). Bafilomycin A1 also induced lysosomal swelling, allowing GFP fluorescence to be visualized within the lysosomal lumen (Fig. 6A). To further examine the luminal localization of PCDH17–GFP, we enlarged endolysosomal compartments by expressing a constitutively active Rab5 mutant (Stenmark et al., 1994; Ohshima et al., 2022). PCDH17–GFP fluorescence was readily detected within the lumen of these enlarged compartments (Fig. 6B). Because the GFP-tagged C terminus of PCDH17 is exposed to the cytosol, its appearance within the lysosomal lumen indicates that PCDH17 is internalized into intraluminal structures through a membrane-remodeling process, potentially involving ESCRT-dependent multivesicular-body biogenesis or microautophagy (Katzmann et al., 2002; Wang et al., 2023). Collectively, these observations indicate that the lysosome-associated pool of PCDH17–GFP is delivered into the lysosomal lumen and subsequently degraded, accounting for the loss of lysosomal PCDH17 fluorescence during starvation.

**Figure 6.**
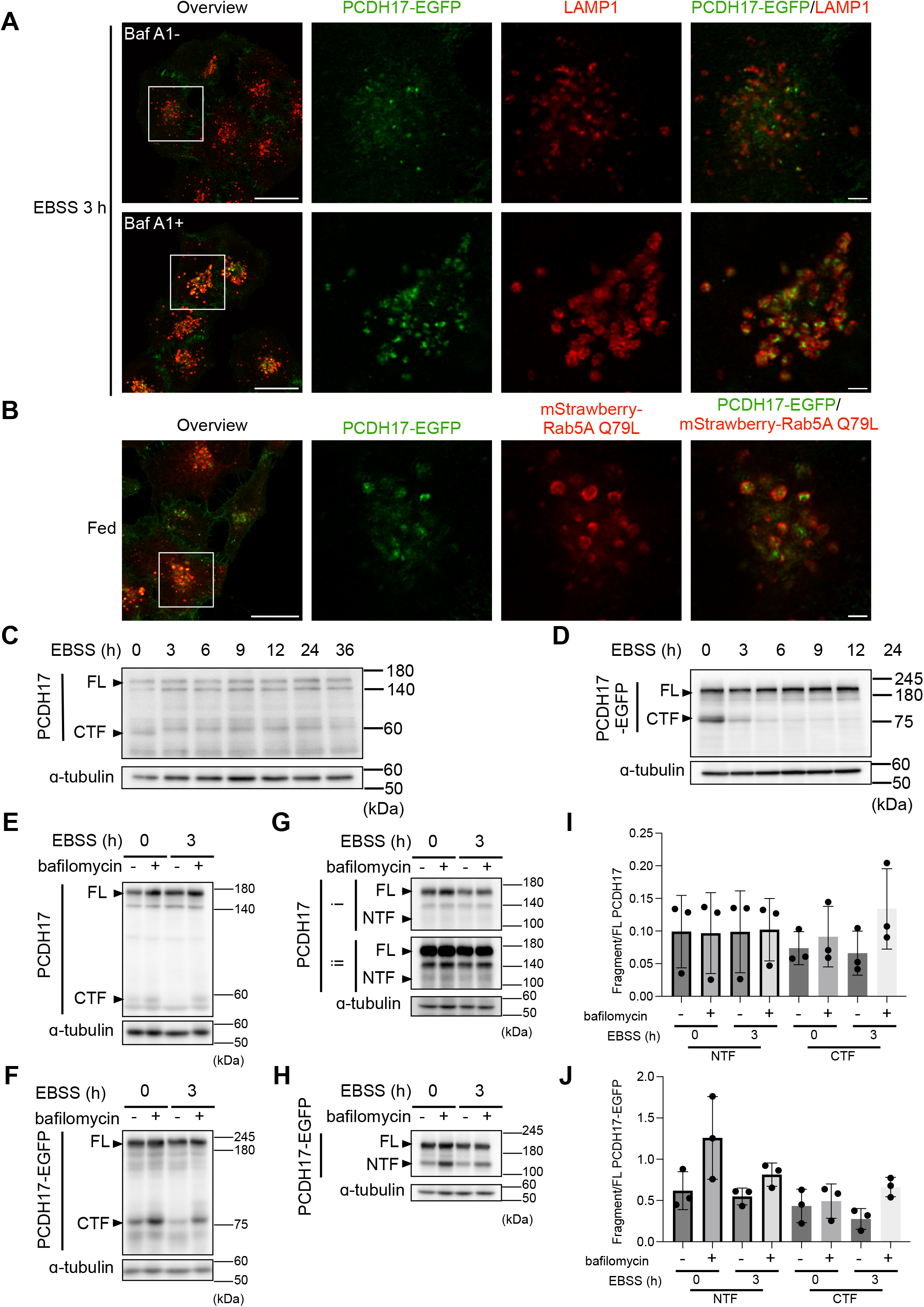
PCDH17 is proteolytically processed, and its C-terminal fragment undergoes enhanced lysosomal degradation during starvation. (A) HeLa cells expressing PCDH17–EGFP were starved in EBSS for 3 h in the presence or absence of 200 nM bafilomycin A1, fixed, stained with anti-Lamp1, and imaged by confocal microscopy. Boxed regions in the overview images are enlarged in the adjacent panels. The enlarged images were deconvolved using Huygens software. Scale bars, 20 μm in the overview images and 2 μm in the enlarged images. (B) HeLa cells coexpressing PCDH17–EGFP and constitutively active mStrawberry–Rab5A Q79L were cultured in complete medium, fixed, and imaged by confocal microscopy. Boxed regions in the overview images are enlarged in the adjacent panels. The enlarged images were deconvolved using Huygens software. Scale bars, 20 μm in the overview images and 2 μm in the enlarged images. (C) HeLa cells were starved in EBSS for 0, 3, 6, 9, 12, 24, or 36 h. Cell lysates were analyzed by immunoblotting using an antibody recognizing the C-terminal region of PCDH17. Full-length PCDH17 and its C-terminal fragment (CTF) are indicated. (D) HeLa cells expressing PCDH17–EGFP were starved in EBSS for 0, 3, 6, 9, 12, or 24 h. Cell lysates were analyzed by immunoblotting with an antibody recognizing the C-terminal region of PCDH17. Full-length PCDH17–EGFP and its CTF are indicated. (E) HeLa cells were maintained in complete medium or starved in EBSS for 3 h in the presence or absence of 200 nM bafilomycin A1. Endogenous full-length PCDH17 and its CTF were detected by immunoblotting with an antibody recognizing the C-terminal region of PCDH17. (F) HeLa cells expressing PCDH17– EGFP were maintained in complete medium or starved in EBSS for 3 h in the presence or absence of 200 nM bafilomycin A1. Full-length PCDH17–EGFP and its CTF were detected by immunoblotting with an antibody recognizing the C-terminal region of PCDH17. (G) HeLa cells were maintained in complete medium or starved in EBSS for 3 h in the presence or absence of 200 nM bafilomycin A1. Cell lysates were analyzed by immunoblotting using an antibody raised against amino acids 18–702 of PCDH17, which recognizes its N-terminal region. Full-length PCDH17 and the N-terminal fragment (NTF) are indicated. (i) and (ii) indicate the immunoblot results with original and adjusted intensities, respectively. (I, J) Quantification of fragment to full length PCDH17 (E, G) and fragment to full length PCDH17-EGFP in (F, H), respectively (n = 3 independent experiments).

To characterize the molecular species undergoing lysosomal degradation, we analyzed HeLa cell lysates by immunoblotting with an anti-PCDH17 antibody recognizing amino acids 812–1159, which encompass the C-terminal cytoplasmic domain. In addition to full-length PCDH17, detected at approximately 164 kDa, this antibody recognized a second species of approximately 56 kDa under nutrient-rich conditions (Fig. 6C). A fragment comprising the transmembrane domain and C-terminal cytoplasmic domain has a predicted molecular mass of approximately 48 kDa. Thus, the 56-kDa species contains the C-terminal cytoplasmic region of PCDH17 and should retain its transmembrane domain. We hereafter refer to this species as the PCDH17 C-terminal fragment (CTF).

During starvation, the endogenous PCDH17 CTF progressively decreased and became undetectable after 3 h (Fig. 6C). Similarly, the CTF derived from PCDH17–EGFP decreased during starvation, whereas the abundance of full-length PCDH17–EGFP remained largely unchanged (Fig. 6D). Bafilomycin A1 prevented the starvation-induced loss of both endogenous and EGFP-tagged CTFs without appreciably affecting the corresponding full-length proteins (Fig. 6E and F). Moreover, bafilomycin A1 increased CTF abundance under both nutrient-rich and starvation conditions (Fig. 6E and F). These results indicate that the CTF undergoes constitutive lysosomal turnover that is enhanced during starvation.

### The N-terminal fragment of PCDH17 constitutes a stable pool within lysosomal compartments

These findings prompted us to investigate the fate of the complementary N-terminal portion of PCDH17. We therefore performed immunoblotting using a second anti-PCDH17 antibody raised against amino acids 18–702, a region located N-terminal to the transmembrane domain. In addition to full-length PCDH17, this antibody detected a species of approximately 112 kDa under nutrient-rich conditions, which we designated the PCDH17 N-terminal fragment (NTF) (Fig. 6G and H).

The region extending from the mature N terminus, following signal-peptide removal, to the beginning of the transmembrane domain has a predicted molecular mass of approximately 77 kDa. The higher apparent molecular mass of the NTF is consistent with glycosylation of the extracellular region, which contains multiple annotated N-linked glycosylation sites (UniProtKB: O14917). Together with the apparent molecular mass of the CTF, these observations suggest that PCDH17 is proteolytically cleaved near its transmembrane domain, generating an NTF composed predominantly of extracellular cadherin repeats.

Notably, the NTF remained readily detectable after 3 h of starvation, whereas the CTF was largely depleted (Fig. 6E–H). Bafilomycin A1 treatment during starvation prevented CTF degradation and markedly increased CTF abundance of endogenous PCDH17 (Fig. 6E). In contrast, the corresponding increase in NTF abundance was comparatively modest (Fig. 6G). In addition, in cells expressing PCDH17-EGFP, the starvation-induced reduction in CTF was more pronounced (Fig. 6F). Although an additional accumulation was unexpectedly detected at the position of the NTF in PCDH17-EGFP-expressing cells, little change in NTF band intensity was observed in the absence of bafilomycin A1 treatment (Fig. 6H). Following bafilomycin A1 treatment, the CTF-to-full-length and NTF-to-full-length ratios converged to similar values of approximately 0.5 (Fig. 6I and J). These results suggest that both fragments undergo lysosomal turnover but that the NTF is substantially more resistant to lysosomal degradation than the CTF. Thus, the NTF constitutes a relatively stable pool of PCDH17 within lysosomal compartments.

### PCDH17 attenuates lysosomal acidification and proteolytic function

Given the sustained autophagic activity observed in PCDH17-depleted cells and the dynamic localization of PCDH17 within lysosomal compartments, we next investigated whether PCDH17 regulates lysosomal homeostasis. We first found that the number of LAMP1-positive lysosomal puncta per cell was significantly increased in PCDH17-knockdown cells relative to cells expressing a scrambled control shRNA. (Fig. 7A and B). Consistent with this observation, PCDH17 depletion increased LysoTracker fluorescence (Fig. 7C and D), which may reflect enhanced lysosomal acidification, expansion of the acidic lysosomal compartment, or both. Importantly, expression of an shRNA-resistant PCDH17 construct rescued the increase in LysoTracker fluorescence, confirming that this phenotype was specifically attributable to PCDH17 depletion (Fig. 7C and D; Fig. S5A).

**Figure 7.**
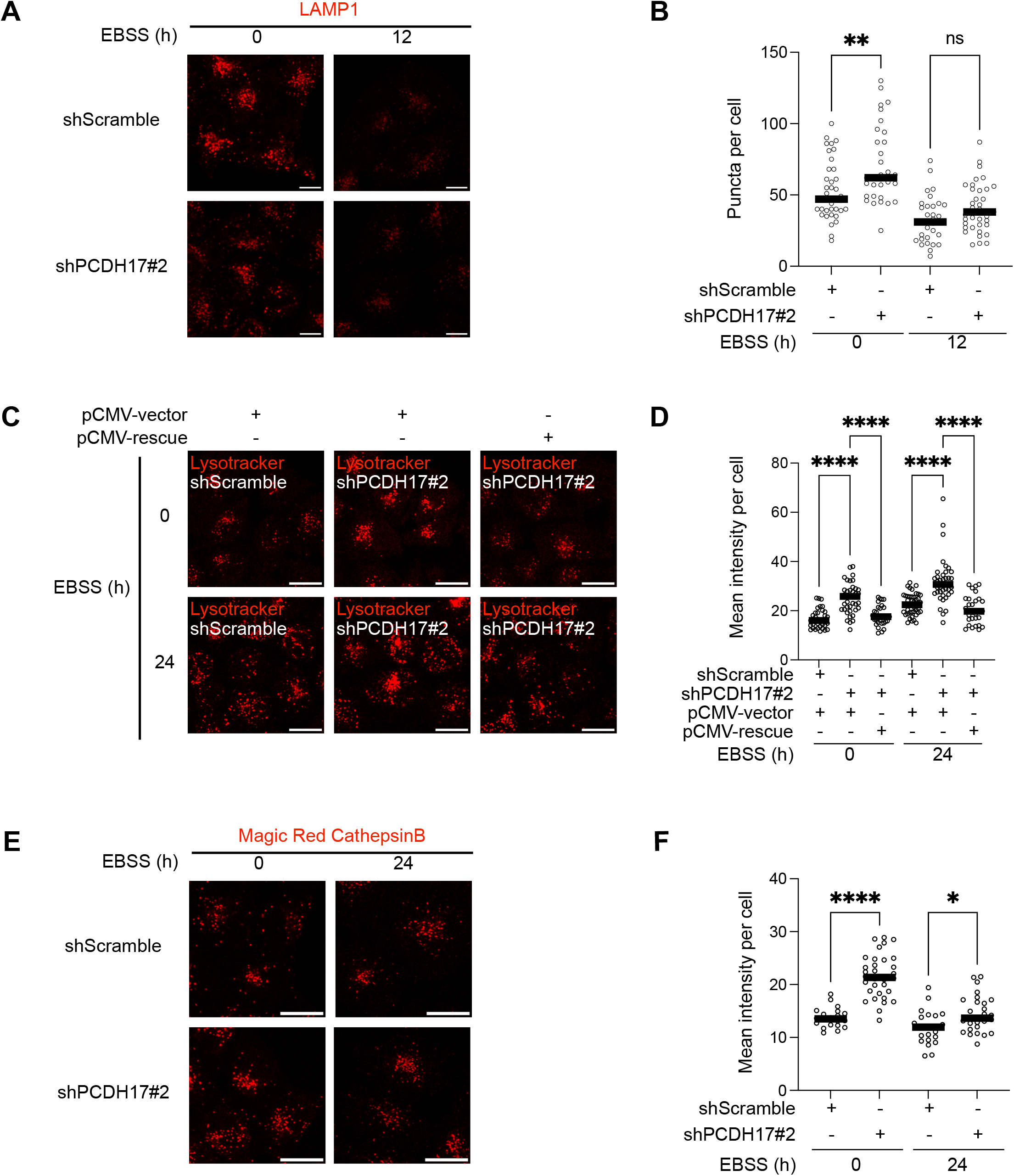
PCDH17 modulates lysosomal abundance, acidification, and proteolytic activity. (A) HeLa cells transduced with shScramble and shPCDH17#2 lentivirus were maintained in complete medium or starved in EBSS for 12 h, fixed, and immunostained for LAMP1. Representative confocal images are shown. Scale bars, 10 μm. (B) Quantification of the number of LAMP1-positive puncta per cell in (A) (n > 30 cells per condition). (C) Control or PCDH17 knockdown Hela cells were transfected with empty pCMV vector or the shPCDH17#2-resistant PCDH17 plasmid (pCMV-rescue) 24 h before starvation as indicated. Cells were maintained in complete medium or starved in EBSS for 24 h, stained with LysoTracker Red DND-99 during the final 30 min of treatment, and imaged live. Representative images are shown. Scale bars, 20 μm. (D) Quantification of LysoTracker fluorescence intensity per cell in (C) (n > 30 cells per condition). (E) Control or PCDH17 knockdown Hela cells were maintained in complete medium or starved in EBSS for 24 h and then subjected to the Magic Red cathepsin B assay. Representative live-cell images are shown. Scale bars, 20 μm. (F) Quantification of Magic Red fluorescence intensity per cell in (E) (n > 20 cells per condition). Statistical significance was assessed by one-way ANOVA (\**p* < 0.05; ** *p* < 0.01; **** *p* < 0.0001).

We next assessed lysosomal proteolytic activity using the Magic Red Cathepsin B assay (Clemente et al., 2022). PCDH17-knockdown cells exhibited significantly higher intracellular cathepsin B activity than scrambled-control cells (Fig. 7E and F), indicating enhanced lysosomal proteolytic function. In addition, the abundance of LAMP1 and mature cathepsin D was moderately increased following PCDH17 depletion (Fig. 8A–C), further supporting increased lysosomal abundance and proteolytic capacity.

**Figure 8.**
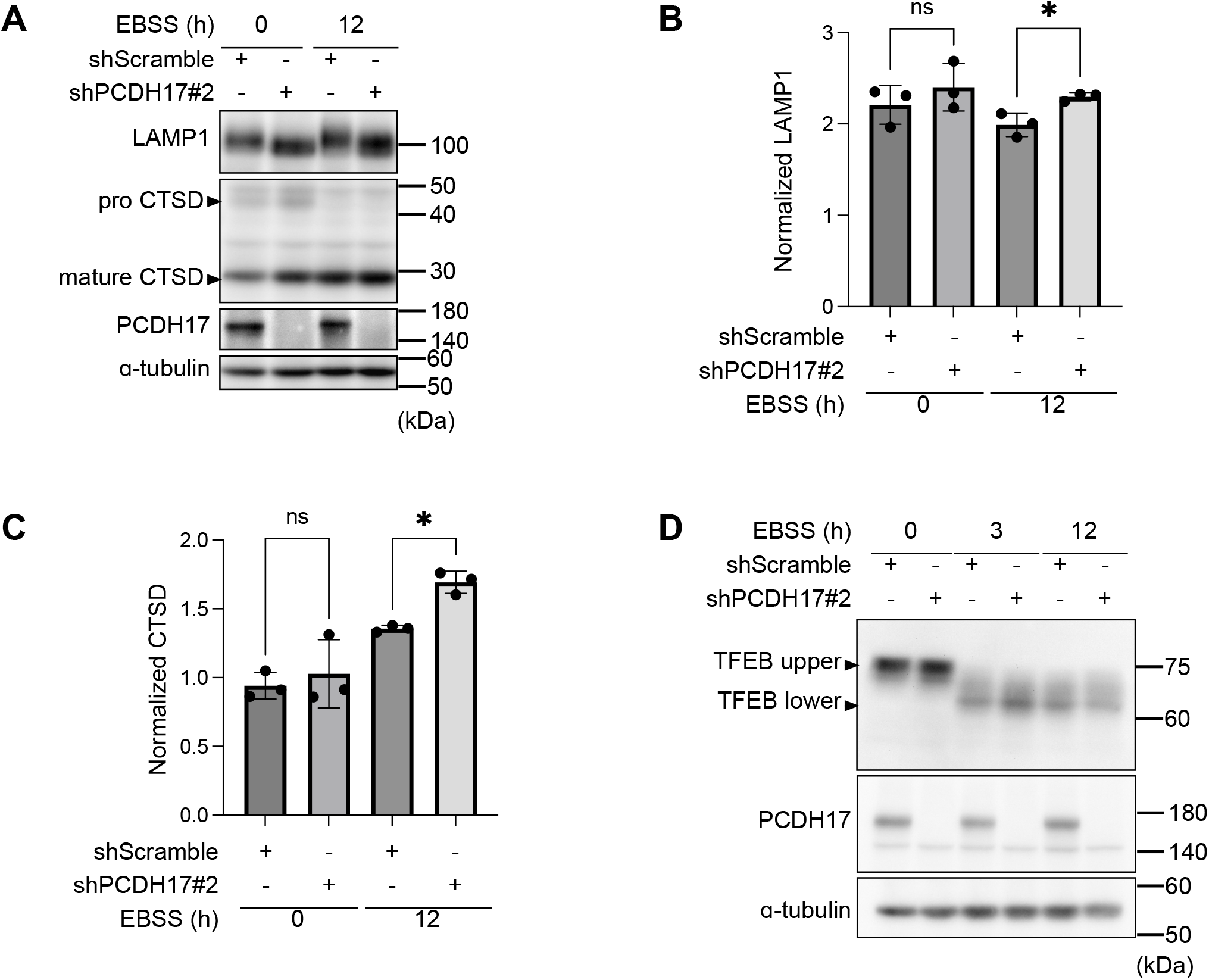
Effects of PCDH17 on TFEB, cathepsin D, and LAMP1 levels. (A and D) Control or PCDH17 knockdown Hela cells were maintained in complete medium or starved in EBSS for 12 h. Cell lysates were analyzed by immunoblotting with antibodies against LAMP1, cathepsin D (A), TFEB (D), PCDH17 (A, D), and the indicated loading control. The precursor and mature forms of cathepsin D are indicated. (B, C) Quantification of LAMP1 (B) and mature cathepsin D (C) abundance in (A), normalized to the loading control (n = 3 independent experiments). Data are presented as the mean ± SD. Statistical significance was assessed by one-way ANOVA (\**p* < 0.05).

TFEB is a principal transcriptional regulator of lysosomal biogenesis and coordinates the expression of lysosome-related genes (Sardiello et al., 2009). Under starvation, it is dephosphorylated and translocated from the cytosol to nucleus, thereby upregulate the lysosome related genes (Roczniak-Ferguson et al., 2012). In PCDH17-knockdown cells, a faster-migrating TFEB species, consistent with a dephosphorylated form of TFEB, was increased under nutrient-rich conditions (Fig. 8D). After 3 h of starvation, this species was readily detected and remained more abundant in PCDH17-knockdown cells than in control cells; however, this difference was no longer apparent after 12 h of starvation; however, this difference was no longer apparent after 12 h of starvation (Fig. 8D).

Conversely, PCDH17 overexpression markedly reduced TFEB protein abundance under nutrient-rich conditions, and this reduction persisted after 3 and 12 h of starvation (Fig. S5B). The electrophoretic mobility of TFEB was also altered, consistent with a shift toward the dephosphorylated form. These changes in TFEB abundance and electrophoretic mobility raised the possibility that PCDH17 modulates lysosomal homeostasis, at least in part, through TFEB. Consistent with this possibility, the number of LAMP1-positive puncta was reduced in PCDH17-overexpressing cells under both nutrient-rich and starvation conditions (Fig. S5C and E). LysoTracker fluorescence was likewise markedly reduced following PCDH17 overexpression (Fig. S5D and F), indicating reduced lysosomal acidification. Consistent with these imaging results, immunoblot analysis revealed reduced amounts of both LAMP1 and mature cathepsin D following PCDH17 overexpression (Fig. S5G–I). Together, these reciprocal loss- and gain-of-function phenotypes indicate that PCDH17 negatively regulates lysosomal abundance, acidification, and proteolytic capacity.

## Discussion

Through an unbiased genome-wide CRISPR knockout screen designed to identify factors involved in the attenuation of autophagy after prolonged starvation, we identified PCDH17 as a regulator of lysosomal function that contributes to the decline in autophagic activity. PCDH17 is a member of the δ-protocadherin subfamily (Redies et al., 2005). Previous studies of protocadherins have focused primarily on their roles in neuronal development and tumorigenesis, as well as their canonical functions in cell–cell adhesion (Redies et al., 2005; Yagi, 2008). Our findings, however, indicate that PCDH17 also performs intracellular functions distinct from its established role at the plasma membrane.

In addition to the well-established plasma membrane pool of PCDH17, we identified a distinct lysosome-associated subpopulation. This finding initially presented an apparent paradox: C-terminally GFP-tagged PCDH17 disappeared from lysosomal compartments during starvation, whereas genetic depletion of PCDH17 impaired the attenuation of autophagy. If lysosomal PCDH17 were eliminated entirely during starvation, its depletion by knockdown or knockout would not be expected to produce an additional phenotype. Resolving this apparent discrepancy was therefore essential for understanding how PCDH17 contributes to the regulation of autophagic activity during prolonged starvation. Our finding that the N-terminal fragment (NTF) of PCDH17 is comparatively resistant to lysosomal degradation provides a potential resolution to this paradox. Specifically, the disappearance of the C-terminal GFP signal does not necessarily indicate the complete loss of lysosomal PCDH17, because a relatively stable NTF persists and may retain the capacity to modulate lysosomal function. PCDH17 also modulates lysosomal degradative activity, potentially through alterations in lysosomal acidification via TFEB regulation (Fig. 7; Fig. 8; Fig. S5 B -I). In addition, the number of LAMP1-positive lysosome increased with PCDH17 depletion (Fig. 7A and B; Fig. S5 C and E). Although the mechanism by which the NTF affects lysosomal activity remains to be defined, its persistence within lysosomal compartments and relative resistance to lysosomal degradation are notable. The NTF consists predominantly of extracellular cadherin repeats, which can mediate Ca^2+^-dependent homophilic interactions in cadherin-family proteins (Brasch et al., 2012). Because lysosomes contain a releasable Ca^2+^ pool (Christensen et al., 2002; Lloyd-Evans et al., 2008), we speculate that such interactions may promote NTF oligomerization within the lysosomal lumen. This higher-order assembly could limit the accessibility of lysosomal proteases and thereby contribute to the relative stability of the NTF.

It has been extensively studied on the endocytic internalization of cell-surface cadherins and their subsequent delivery to lysosomes for degradation (Cadwell et al., 2016; Delva and Kowalczyk, 2009). These processes regulate the abundance of cadherins at the plasma membrane and thereby modulate cell–cell adhesion. By contrast, our findings indicate that the lysosome-associated pool of PCDH17 is delivered through the biosynthetic ER–Golgi pathway rather than through endocytosis from the plasma membrane (Fig. 5). This observation suggests that newly synthesized PCDH17 is sorted at, or shortly after passage through, the Golgi apparatus into distinct trafficking routes directed toward either the plasma membrane or lysosomal compartments. A conceptually similar nutrient-responsive biosynthetic sorting decision has been described for the yeast general amino acid permease Gap1. Under poor nitrogen conditions, newly synthesized Gap1 is delivered to the plasma membrane, whereas loss of Npr1 or growth under preferred nitrogen conditions diverts newly synthesized Gap1 from the Golgi to the vacuole without prior delivery to the cell surface (De Craene et al., 2001). Thus, regulated post-Golgi sorting provides a plausible model for the partitioning of newly synthesized PCDH17. However, the molecular machinery involved and whether PCDH17 reaches lysosomes directly or via endosomal intermediates remain to be determined. Nevertheless, the direct biosynthetic delivery of PCDH17 to lysosomes suggests that lysosomal localization represents a functionally relevant destination rather than merely the terminal disposal of cell-surface PCDH17. Because the GFP-tagged C terminus of PCDH17 is oriented toward the cytosol, its detection within the lysosomal lumen suggests that PCDH17 is incorporated into intraluminal membrane structures. Similar intraluminal localization and trafficking to multivesicular endolysosomal compartments have been reported for γ-protocadherins (Hanson et al., 2010; O’Leary et al., 2011; Shonubi et al., 2015). This process may involve ESCRT-dependent internalization, a conserved pathway for the selective delivery of lysosomal membrane proteins into the lumen (Zhang et al., 2021).

At the outset of this study, we aimed to identify factors responsible for the attenuation of autophagy during prolonged starvation. Given that nutrient-responsive regulators of autophagy, most notably the mTORC1–ULK1 signaling axis, have been well characterized (Kim et al., 2011), we initially anticipated that additional components of analogous signaling pathways might underlie this attenuation. Rather than implicating an upstream nutrient-sensing pathway, however, our findings unexpectedly pointed to regulation of lysosomal degradative capacity—the terminal stage of the autophagic process. These findings emphasize that overall autophagic flux is determined not only by upstream signaling and the efficiency of autophagosome biogenesis but also by the capacity of lysosomes to degrade autophagic cargo. Indeed, lysosomal adaptation to starvation is itself temporally regulated. TFEB, a master regulator of lysosomal biogenesis and function, undergoes rapid activation and nuclear translocation upon nutrient deprivation (Settembre et al., 2012), followed by partial adaptation rather than a sustained monotonic increase during continued starvation (Ruolo et al., 2025). Nutrient deprivation typically suppresses mTORC1 activity, leading to dephosphorylation and nuclear translocation of TFEB. Under nutrient-rich conditions, mTORC1 phosphorylates TFEB at Ser211, promoting its cytoplasmic retention (Roczniak-Ferguson et al., 2012). Upon mTORC1 inhibition, calcineurin dephosphorylates TFEB, allowing TFEB to translocate to the nucleus and activate the expression of lysosomal and autophagy-related genes (Ruolo et al., 2025). Interestingly, even under nutrient-rich conditions, dephosphorylated TFEB was increased in PCDH17 knockdown cells (Fig. 8D), and a marked reduction of both hyper phosphorylated and dephosphorylated TFEB was also observed in PCDH17 overexpression (Fig. S5B), raising the possibility that PCDH17 modulates lysosomal activity through a TFEB-dependent mechanism. Given that the lysosomal NTF may affect the lysosomal Ca²⁺ pool, the Ca²⁺-dependent phosphatase calcineurin represents a potential link between PCDH17 and TFEB regulation at the lysosome. Thus, PCDH17 may influence TFEB phosphorylation status through modulation of lysosomal Ca²⁺-dependent calcineurin activity. In parallel, mTOR activity can recover during prolonged starvation through the generation of autolysosomal degradation products, thereby attenuating autophagy and promoting autophagic lysosome reformation (Yu et al., 2010). Thus, changes in autophagic activity during prolonged starvation cannot be understood solely in terms of the pathways governing autophagy induction.

Emerging evidence suggests that protocadherins participate in intracellular processes beyond cell– cell adhesion (Phillips et al., 2017). For example, expression of Pcdh-γA3 or Pcdh-γB2 induces the formation of juxtanuclear membrane tubules that recruit lipidated LC3 but are distinct from conventional autophagic structures. Correlative light and electron microscopy, together with colocalization with the lysosomal marker LAMP2, indicated that these tubules arise from lysosomal compartments (Hanson et al., 2010). Subsequent studies identified a motif within the variable cytoplasmic domain that is required for γ-protocadherin trafficking through the endolysosomal system and for the associated membrane tubulation (O’Leary et al., 2011; Shonubi et al., 2015). These observations suggest that protocadherins can participate in intracellular membrane trafficking and remodeling independently of their canonical adhesive functions.

Although PCDH17 is preferentially expressed in the nervous system and has established roles in neuronal development and circuit formation (Hoshina et al., 2013; Hayashi et al., 2014; Asakawa and Kawakami, 2018), its expression is not restricted to neural tissues; notably, we detected endogenous PCDH17 in HeLa cells. This broader expression pattern may be pathophysiologically relevant. PCDH17 promoter hypermethylation and reduced expression have been reported in several epithelial cancers, including gastric, colorectal, and esophageal squamous-cell carcinomas, where PCDH17 has been proposed to exert tumor-suppressive functions (Haruki et al., 2010; Hu et al., 2013). In colorectal-cancer models, PCDH17 expression has also been reported to influence sensitivity to 5-fluorouracil (Liu et al., 2019), and genetic variation at the *PCDH17* locus has been associated with susceptibility to major mood disorders (Chang et al., 2018). These observations raise the possibility that PCDH17-dependent regulation of lysosomal function may influence cellular adaptation to chronic metabolic stress in both physiological and disease contexts. Future studies should define how PCDH17 proteolytic processing and its lysosome-associated N-terminal fragment regulate lysosomal acidification and proteolysis, and should test whether this pathway affects cell survival, tumor progression, therapeutic responses, or neuronal function in disease-relevant models.

## Materials and methods

### Cell culture, transfection, transduction

HeLa cells were obtained from laboratory stocks. HeLa cells and their derivatives were maintained in high-glucose Dulbecco’s modified Eagle’s medium (DMEM; Sigma-Aldrich, D6429) supplemented with 10% fetal bovine serum (FBS; Invitrogen) at 37°C in a humidified atmosphere containing 5% CO₂. For nutrient-starvation experiments, the culture medium was replaced with Earle’s balanced salt solution (EBSS; Sigma-Aldrich, E2888) for the indicated durations. Plasmid transfections were performed using Lipofectamine 2000 (Invitrogen, 11668019).

For retrovirus production, Plat-E packaging cells (Morita et al., 2000) were transfected with the indicated retroviral plasmids using polyethylenimine (PEI) (Polysciences, 23966). At 24 h after transfection, the transfection medium was replaced with fresh growth medium. Virus-containing supernatants were collected at 48 and 72 h after transfection and passed through a 0.45-μm filter (Millipore, SLHVR33RS). Target cells were seeded 24 h before transduction and incubated with the filtered viral supernatants supplemented with 10 μg/mL polybrene (Sigma-Aldrich, H9268-5G). After 24 h, the viral medium was replaced with fresh growth medium. Following an additional 24-h recovery period, transduced cells were selected with either 2 μg/mL puromycin or 5 μg/mL blasticidin, as appropriate.

For the production of shRNA-expressing lentiviruses, HEK293T cells were cotransfected with the packaging plasmid psPAX2 (Addgene plasmid #12260), the envelope plasmid pMDG (VSVG) (Sušac et al., 2022), and the appropriate lentiviral transfer plasmid using PEI. Subsequent collection and filtration of virus-containing supernatants, transduction of target cells, and antibiotic selection were performed as described above for retrovirus production.

For transient PCDH17 depletion, cells were transfected with 100 nM control siRNA (siControl; Sigma-Aldrich, SIC001) or one of two siRNAs targeting PCDH17—siPCDH17 #1 (Sigma-Aldrich, SASI_Hs01_00138036) and siPCDH17 #2 (Sigma-Aldrich, SASI_Hs01_00138038)— using Lipofectamine RNAiMAX Transfection Reagent (Invitrogen, 13778150) according to the manufacturer’s instructions. Cells were analyzed 24 h after transfection.

### Plasmids

pMRX-IP-HaloTag7-LC3 (Addgene plasmid #184899) and pMRX-IB-HaloTag7-mGFP (Addgene plasmid #184903) were obtained from Addgene and have been described previously (Yim et al., 2022). pMRX-IB-HaloTag7-LC3 was generated by replacing the HaloTag7-mGFP coding sequence in pMRX-IB-HaloTag7-mGFP with the HaloTag7-LC3 coding sequence. These plasmids were used for retrovirus production and subsequent generation of cells stably expressing the indicated HaloTag7 fusion proteins for HaloTag-based degradation assays (Yim et al., 2022). All shRNA constructs were purchased from Sigma-Aldrich; their sequences and product information are provided in Table S2. pCMV-SPORT6-PCDH17 (RIKEN, HGX025117) was used as the source of the human PCDH17 coding sequence. To generate PCDH17–EGFP, the PCDH17 coding sequence was cloned into the pMRX-IP vector with EGFP fused in frame to its C terminus. The corresponding empty pCMV vector, lacking the PCDH17 coding sequence, was used as a control. For rescue experiments, a shRNA-resistant PCDH17 construct (pCMV-rescue) was generated by introducing synonymous substitutions into the shPCDH17 #2 target site. Specifically, the original target sequence, 5′-CGAGACACAAGACGAGTACAA-3′, was changed to 5′-GGAAACTCAAGACGAGTACAA-3′. pMRX-IB LAMP1-RFP is a laboratory stock (Lu et al., 2025). pMRX-IP GFP-ATG5 plasmid (Ikari et al., 2020) was a gift from Dr. Akiko Kuma, The University of Osaka. mStr–Rab5a-QL plasmid (Hiragi et al., 2022; Ohbayashi et al., 2012) was a gift from Dr. Fukuda Mitsunori, Tohoku University.

### Antibodies

The following primary antibodies were used for immunofluorescence: rabbit anti-calreticulin (1:300; Proteintech, 27298-1-AP), rabbit anti-LC3 (1:1,000; Medical & Biological Laboratories, PM036), mouse anti-GM130 (1:200; BD Biosciences, 610822), rabbit anti-GM130 (1:1,000; Cell Signaling Technology, 12480), rabbit anti-EEA1 (1:100; Cell Signaling Technology, 2411), and mouse anti-LAMP1 (1:500; Santa Cruz Biotechnology, H4A3). The secondary antibodies were Alexa Fluor 568–conjugated goat anti-rabbit IgG (1:1,000; Thermo Fisher Scientific, A10042), Alexa Fluor 568–conjugated goat anti-mouse IgG (1:1,000; Thermo Fisher Scientific, A10037), and Alexa Fluor 647–conjugated goat anti-rabbit IgG (1:1,000; Thermo Fisher Scientific, A31573).

For immunoblotting, the following primary antibodies were used: mouse anti-HaloTag (1:1,000; Promega, G21A), rabbit anti-phospho-p70 S6 kinase (1:2,000; Cell Signaling Technology, 9234), rabbit anti-p70 S6 kinase (1:2,000; Cell Signaling Technology, 2708), rabbit anti-phospho-ULK1 (1:1,000; Cell Signaling Technology, 6888), rabbit anti-ULK1 (1:1,000; Cell Signaling Technology, 8054), rabbit anti-PCDH17 (C-terminal immunogen) (1:1,000; Proteintech, 17915-1-AP), sheep anti-PCDH17 (N-terminal immunogen) (1:1,000; R&D systems, AF7669), goat anti-cathepsin D (1:1,000; Santa Cruz Biotechnology, C-20), rabbit anti-TFEB (1:1000, Cell Signaling Technology, 4240), mouse anti-β-actin (1:20,000; Proteintech, 66009-1-Ig), mouse anti-α-tubulin (1:20,000; Proteintech, 66031-1-Ig). The secondary antibodies were horseradish peroxidase (HRP)– conjugated goat anti-mouse IgG (1:10,000; Jackson ImmunoResearch, 115-035-003) and HRP-conjugated goat anti-rabbit IgG (1:10,000; Jackson ImmunoResearch, 111-035-003), HRP-conjugated goat anti-sheep IgG (1:10,000; Jackson ImmunoResearch, 713-035-147), Rabbit Anti-Goat IgG H&L (HRP) (1:10,000; Abcam, AB6741).

### Reagents

The following reagents were used: bafilomycin A1 (BioViotica, BVT-0252-M001), wortmannin (FUJIFILM Wako, 230-02341), dynasore (AdipoGen, AG-CR1-0045), brefeldin A (Cell Signaling Technology, 9972S), cycloheximide (FUJIFILM Wako, 037-20991), digitonin (Millipore, 300410-250MG), puromycin (FUJIFILM Wako, 160-23151), blasticidin (KNF, KK-400), and ProLong Glass Antifade Mountant (Invitrogen, P36984).

### Immunoblotting

Cells were seeded and treated as indicated for each experiment and then lysed on ice for 20 min in lysis buffer containing 5 mM Tris-HCl (pH 7.4), 150 mM NaCl, and 1% Triton X-100, supplemented with a protease inhibitor cocktail (Roche, 11873580001). Cell lysates were clarified by centrifugation (16,500 × g, 10 min, 4°C), and the resulting supernatants were mixed with 5 × sample buffer containing 250 mM Tris-HCl, 8% SDS, 0.1% bromophenol blue, 40% glycerol, and 100 mM dithiothreitol. Samples were heated at 70°C for 10 min, resolved by SDS–PAGE, and transferred onto polyvinylidene difluoride membranes (PVDF; Millipore, IPVH00010). The membranes were blocked with 5% skim milk in phosphate-buffered saline-Tween 20 (PBS-T) (137 mM NaCl, 2.7 mM KCl, 10 mM Na₂HPO4, 1.8 mM KH₂PO₄ and 0.1% Tween 20) for 1 h at room temperature and then sequentially incubated with the indicated primary antibody diluted in 1% skim milk in PBS-T and secondary antibodies diluted in PBS-T for 1 h each at room temperature.

### Immunofluorescence

Cells were seeded on coverslips (Matsunami, CS01813) and treated as indicated for each experiment. Cells were fixed with 4% paraformaldehyde (PFA; FUJIFILM Wako, 163-20145) for 20 min at room temperature and permeabilized with 50 μg/mL digitonin in PBS for 10 min at room temperature. Samples were then blocked with 0.2% gelatin (FUJIFILM Wako, 076-02765) in PBS for 1 h at room temperature. Primary and secondary antibodies were diluted in 0.2% gelatin in PBS, and samples were sequentially incubated with each antibody for 1 h at room temperature. Finally, the samples were mounted using ProLong Glass Antifade Mountant (Invitrogen, P36984) for imaging.

### Lentiviral CRISPR library preparation and screening

The human Brunello sgRNA library in lentiCRISPR v2 (Addgene #73179) was amplified in DH5α competent cells (SMOBIO, CC5202) as described previously (Doench et al., 2016). Library representation was confirmed by NovaSeq X Plus sequencing. HeLa cells stably expressing HaloTag7–LC3 (3 × 10⁸ cells distributed across fifteen 150-mm culture dishes) were transduced with the lentiviral library at a transduction efficiency of approximately 10%. At 24 h after transduction, cells were selected with 2 μg/mL puromycin for 7 days.

For each screen, 3 × 10⁷ puromycin-selected cells were starved in EBSS for 24 h, with 200 nM bafilomycin A1 added during the final 1 h of starvation. Cells were then subjected to the Takahashi assay and sorted using a FACSAria Fusion cell sorter (BD Biosciences). Debris was excluded on the basis of forward- and side-scatter profiles. MIL and MPL fluorescence signals were acquired in the Alexa Fluor 488 and PE channels, respectively. The autophagy-positive sorting gate was positioned outside the fluorescence distribution of parental, library-untransduced control cells processed under the same conditions and was used to collect cells with elevated MPL fluorescence relative to MIL fluorescence.

In parallel, an aliquot of the puromycin-selected library-transduced cells was collected before starvation and was not exposed to bafilomycin A1, the Takahashi assay, or flow-cytometric sorting. This untreated library pool served as the reference population for subsequent sgRNA-enrichment analysis. The sorted and reference populations were processed for genomic DNA extraction and sgRNA sequencing as described below. Three independent screens were performed.

Genomic DNA was extracted from the reference and sorted cell populations using phenol:chloroform alcohol (25:24:1; PCI; NIPPON GENE, 311-90151). To maintain adequate representation of the sgRNA library, a total of 40 μg of genomic DNA from each population was distributed across 4 parallel PCR reactions, with 10 μg of genomic DNA used per 100-μL reaction. Each reaction contained KOD One PCR Master Mix (TOYOBO, KMM-101) and 0.5 μM each of the P5 and barcoded P7 primers. The sgRNA-containing regions were amplified while simultaneously incorporating Illumina sequencing adapters and a sample-specific eight-nucleotide index. The primer sequences were as follows: P5, 5′-AATGATACGGCGACCACCGATCTACACTCTTTCCCTACACGACGCTCTTCCGATCTTTG TGGAAAGGACGAAACACCG-3′; and barcoded P7, 5′-CAAGCAGAAGACGGCATACGAGATNNNNNNNNGTGACTGGAGTTCAGACGTGTGCTCTTCCGATCTCCAATTCCCACTCCTTTCAAGACCT-3′, where NNNNNNNN represents the sample-specific index sequence.

PCR amplification was performed using the following program: initial denaturation at 95°C for 1 min; 35 cycles of denaturation at 95°C for 30 s, annealing at 53°C for 30 s, and extension at 68°C for 1 s; followed by a final extension at 68°C for 10 min. Parallel reactions from each sample were pooled, and the amplified products were resolved by electrophoresis on a 2% agarose gel. Bands of the expected size (285 bp) were excised and purified using the QIAquick Gel Extraction Kit (QIAGEN, 28704). Libraries were normalized to equimolar concentrations, pooled, and sequenced using paired-end 100-bp reads on a NovaSeq X Plus platform at the NGS Core Facility, Research Institute for Microbial Diseases (RIMD), The University of Osaka.

### Sequencing data analysis

Sequencing reads were processed using count_spacers.py, a Python script described by Joung et al. (Joung et al., 2017), to identify sgRNA sequences and generate read counts for each sgRNA. Read counts were normalized to total mapped reads within each sample. Enrichment of individual sgRNAs was calculated as the log₂ fold change in normalized abundance in the sorted population relative to the reference population, filtering out sgRNAs with zero reads in either population. sgRNAs were assigned to their corresponding target genes according to the library annotation. Gene-level candidates were prioritized based on the recurrence of gene symbols across the three independent screens. The complete sgRNA-level dataset, gene-level hit list, and criteria used for candidate refinement are provided in Table S1.

### HaloTag-based autophagic degradation assay using pulse–chase ligand labeling

HeLa cells stably expressing HaloTag7-mGFP were seeded at 5 × 10^5^ cells per well in 6-well plate and cultured for 16 h before labeling. The cells were pulse-labeled by incubation with 100 nM tetramethylrhodamine (TMR) HaloTag ligand (Promega, G8251), diluted in complete growth medium, for 20 min at 37°C. The labeling medium was then removed, and the cells were washed once with prewarmed ligand-free DMEM or EBSS to remove unbound ligand. For the chase period, cells were incubated in the corresponding ligand-free medium—DMEM for nutrient-rich conditions or EBSS for starvation conditions—for the indicated durations.

Cells were collected at the indicated time points, washed with ice-cold PBS, and lysed in lysis buffer mentioned above. Protein concentrations were determined using Bradford Protein Assay (Bio-rad, 5000006) (Bradford, 1976), and equal amounts of protein were separated by SDS–PAGE. TMR-labeled HaloTag7 proteins were detected by Gene Gnome-5 chemiluminescence detector (Syngene). The abundance of the TMR-labeled species was quantified using ImageJ and normalized to the combined intensity of the HaloTag7-mGFP and HaloTag7 bands.

### Takahashi assay

The Takahashi assay (Takahashi et al., 2019, 2018) was performed by sequential labeling of HaloTag7-LC3 with a membrane-impermeant Alexa Fluor 488 HaloTag ligand (MIL) and a membrane-permeant TMR HaloTag ligand (MPL).

Flow cytometric analysis. HeLa cells stably expressing HaloTag7-LC3 were seeded at 1 × 10^6^ cells per 60-mm dish and cultured under the indicated nutrient conditions. Cells were detached with 1 × trypsin (FUJIFILM Wako, 208-17251), neutralized with complete growth medium, collected by centrifugation at 250 × g for 3 min, and washed once with PBS. Approximately 2 × 10^6^ cells per sample were resuspended in 400 μL of 1× mitochondrial assay solution (MAS) containing 220 mM mannitol, 70 mM sucrose, 10 mM KH₂PO₄, 5 mM MgCl₂, 2 mM HEPES, and 1 mM EGTA, adjusted to pH 7.4 by NaOH. The plasma membrane was selectively permeabilized, and accessible HaloTag7-LC3 was simultaneously labeled by incubation in MAS containing 10 μg/mL digitonin and Alexa Fluor 488 HaloTag ligand (Promega, G1001; 1:1,000 dilution; final concentration, 1 μM) for 15 min at 37°C. Samples were protected from light during all labeling steps.

Following MIL labeling, cells were rinsed with 1 mL PBS, collected by centrifugation at 250 × g for 3 min. Cells were then resuspended in 400 μL Hanks’ balanced salt solution (HBSS; Gibco, 14175095) supplemented with 2% FBS and TMR HaloTag ligand (Promega, G8251; 1:2,500 dilution; final concentration, 2 μM) and incubated for 30 min at 37°C. After labeling, cells were washed three times with HBSS containing 2% FBS and resuspended in 1 mL of the same buffer for flow cytometry.

Samples were analyzed using a BD FACSAria Fusion flow cytometer (BD Biosciences). Debris was excluded on the basis of forward- and side-scatter profiles, and single cells were selected using FSC-A versus FSC-W gating. Alexa Fluor 488 and TMR fluorescence were detected using the AF488 and PE channels, respectively. Unlabeled cells and cells labeled individually with each ligand were used to establish background fluorescence and compensation settings. At least 100,000 single-cell events were acquired per sample. Data were analyzed using floreada.io, and autophagic activity was quantified as TMR mean fluorescence intensity.

Fluorescence imaging. HeLa cells stably expressing HaloTag7-LC3 were seeded at 5 × 10^4^ cells per well onto 12-mm coverslips and cultured for 16 h before treatment. Following incubation under the indicated nutrient conditions, cells were washed once with prewarmed PBS and incubated in 1× MAS containing 10 μg/mL digitonin and Alexa Fluor 488 HaloTag ligand (Promega, G1001; 1:1,000 dilution; final concentration, 1 μM) for 15 min at 37°C. Cells were then fixed with 4% paraformaldehyde in PBS for 5 min at 37°C. After three washes with PBS, fixed cells were incubated with TMR HaloTag ligand (Promega, G8251; 1:2,500 dilution; final concentration, 2 μM) in PBS for 30 min at 37°C. Cells were subsequently washed three times with PBS, and coverslips were mounted using ProLong Glass Antifade Mountant (Thermo Fisher Scientific, P36984).

Images were acquired using a Leica TCS SP8 confocal microscope equipped with a HC PL APO CS2 63×/1.40 numerical aperture oil-immersion objective. Identical laser power, detector gain, pinhole, and acquisition settings were used for all samples within each experiment. Alexa Fluor 488 and TMR signals were collected sequentially to minimize spectral bleed-through. Images were analyzed using ImageJ/Fiji, and TMR fluorescence intensity per cell was quantified after background subtraction. At least 30 cells were analyzed per condition in each independent experiment.

### LysoTracker assay

Cells were seeded at 0.5 × 10^6^ cells per dish] in glass-bottom dishes (Matsunami Glass, D11350H) 16 h before imaging. At the time of the assay, cells were approximately 60-80% confluent. For nutrient-rich conditions, cells were maintained in DMEM; for starvation conditions, the culture medium was replaced with EBSS for the indicated duration. LysoTracker Red DND-99 (Invitrogen, L7528) was added to the corresponding medium at a final concentration of 50 nM during the final 30 min of each treatment. Cells were incubated with LysoTracker for 30 min at 37°C, protected from light, and then washed once with prewarmed dye-free DMEM or EBSS, as appropriate. Cells were immediately imaged in the corresponding dye-free medium using a Leica TCS SP8 microscope live-cell imaging system equipped with a HC PL APO CS2 63×/1.40 numerical aperture oil-immersion objective. Temperature was maintained at 37°C during image acquisition.

LysoTracker fluorescence was excited at 561 nm, and emission was collected over 590-650 nm. Laser power, exposure time, detector gain, pinhole, and all other acquisition settings were kept identical among samples within each experiment. At least 6 randomly selected fields and 20 cells were imaged per condition in each independent experiment. Images were analyzed using ImageJ/Fiji. Cell boundaries were defined using transmitted-light images. LysoTracker staining was quantified as mean fluorescence intensity per cell

### Magic Red Cathepsin B assay

Intracellular cathepsin B activity was measured using the Magic Red Cathepsin B Assay Kit (ImmunoChemistry Technologies, 937). Cells were seeded at 0.5 × 10^6^ cells per dish in glass-bottom dishes 24 h before the assay and subjected to the indicated experimental conditions. The Magic Red cathepsin B substrate was freshly prepared as a 1× working solution in DMEM or EBSS according to the manufacturer’s instructions. Where cells underwent a longer treatment, the substrate was added during the final 30 min of the treatment. Cells were incubated with the substrate for 30 min at 37°C in the dark. The substrate-containing medium was then removed, and the cells were rinsed twice with prewarmed dye-free DMEM or EBSS and immediately imaged in DMEM or EBSS.

Live-cell images were acquired using a Leica TCS SP8 microscope imaging system equipped with a HC PL APO CS2 63×/1.40 numerical aperture oil-immersion objective. Temperature was maintained at 37°C during image acquisition. Magic Red fluorescence was excited at 561 nm, and emission was collected over 610-640 nm. Laser power, exposure time, detector gain, and all other acquisition settings were kept identical among samples within each experiment. At least 6 randomly selected fields and 20 cells were analyzed per condition in each independent experiment.

Images were analyzed using ImageJ/Fiji. Cell boundaries were defined using transmitted-light images. Cathepsin B activity was quantified as mean fluorescence intensity per cell

### Imaging and image analysis

Fixed and live-cell samples were imaged using a Leica TCS SP8 confocal laser-scanning microscope (Leica Microsystems) equipped with an HC PL APO CS2 63×/1.40 numerical aperture oil-immersion objective. Images were acquired using Leica Application Suite X software (version 2.0.1.14392). Fluorophores were excited using the appropriate 488-, 561-, 633-nm laser lines, and fluorescence was detected over the corresponding emission ranges using HyD and PMT detectors.

When multiple fluorophores were imaged, channels were acquired sequentially to minimize spectral bleed-through. Images were collected at 1024 × 1024-pixel resolution with a pixel size of 0.18 μm, a scan speed of 200 Hz, and a pinhole diameter of 1.0 Airy unit. Representative images are presented as single optical sections, as indicated in the figure legends. Laser power, detector gain, offset, pinhole size, and all other acquisition settings were kept identical among samples compared within each experiment. Images were acquired below detector saturation.

Fluorescence intensity and puncta number were quantified using ImageJ/Fiji (National Institutes of Health; version 1.54p). Cell boundaries were defined using transmitted-light images. Fluorescence intensity was quantified as mean fluorescence intensity per cell. For puncta analysis, a uniform intensity threshold was applied to all images within an experiment. Objects with an area of 0-Infinity μm² and a circularity of 0–1 were defined as puncta and counted using the Analyze Particles function. Puncta intersecting the cell boundary were included. Where indicated, colocalization was quantified using Pearson’s correlation coefficient with the Coloc 2 ImageJ plugin. At least 6 randomly selected fields and 30 cells were analyzed per condition in each of 2 independent experiments.

Selected image stacks were deconvolved using Huygens essential software (version 15.10.0p5 64b). Unless otherwise indicated, deconvolution was used only to improve the presentation of representative images; quantitative analyses were performed using the original, non-deconvolved images.

For live-cell imaging, cells were seeded in 35-mm glass-bottom dishes (Matsunami Glass, D11350H) 16 h before observation and maintained in the indicated imaging medium. Imaging was performed using the same Leica TCS SP8 microscope equipped with a stage-top incubation chamber maintained at 37°C. For time-lapse experiments, images were acquired every 2.58 s for 28.38 s. Imaging parameters were kept constant across experimental conditions, and laser exposure was minimized to reduce photobleaching and phototoxicity.

### Statistics

Statistical analyses were performed using GraphPad Prism 8 (GraphPad Software). Error bars in all figures indicate standard deviation (SD). Statistical significance was assessed by one-way analysis of variance (ANOVA) or Kruskal-Wallis test as indicated in the figure legends. Significance levels are indicated as follows: \*\*\*\**p* < 0.0001, \*\*\**p* < 0.001, \*\**p* < 0.01, and \**p* < 0.05.

## Acknowledgement

This study was funded by Grants-in-Aid for Scientific Research KAKENHI (20H05326, 22H04647, 23H02475, 26K01963 、26H01630) to T.N. and (20K15789, 25K09629), Hirose Foundation to S-L.L, and JST SPRING (JPMJSP2138) to B.C. We acknowledge the NGS core facility at the Research Institute for Microbial Diseases of The University of Osaka for the sequencing and data analysis. We thank Dr. Eiji Morita at Hirosaki University for providing pMDG. We thank Dr. Akiko Kuma at The University of Osaka for providing pMRX-IP GFP-ATG5. We thank Dr. Mitsunori Fukuda at Tohoku University for providing mStr-Rab5a QL.

## Author contributions

T.N. and B.C. conceived and designed the study. B.C. performed all experiments, analyzed the data, and wrote the original draft of the manuscript. S-L.L. contributed to scientific discussions and provided experimental advice. T.N. supervised the project, and revised and edited the manuscript. All authors discussed the results and approved the final manuscript.

## Supplementary figure, table and video legends

**Figure S1.**
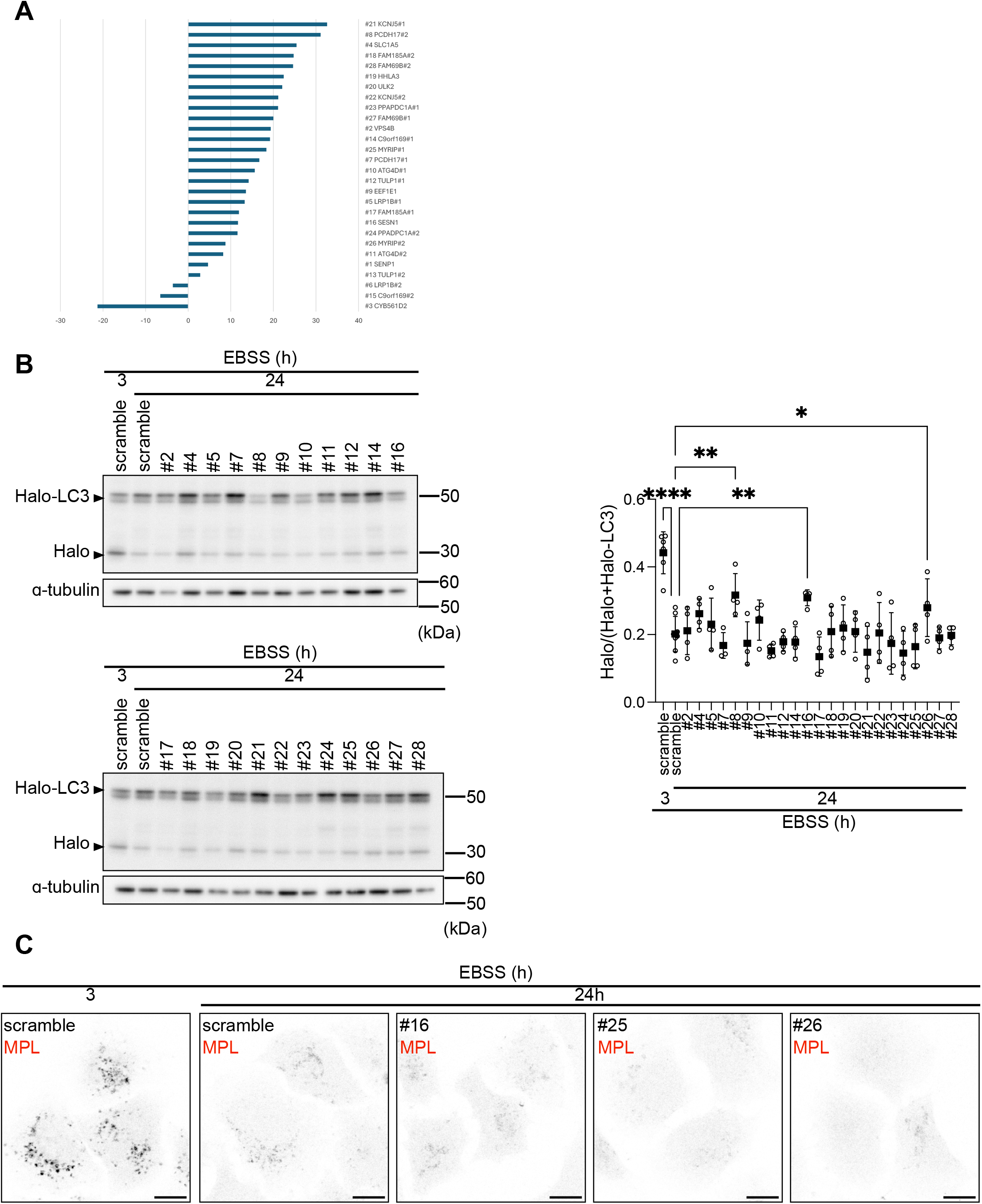
Secondary screening identifies PCDH17 as a regulator of autophagy attenuation. (A) HeLa cells expressing HaloTag7–LC3 were transduced lentivirus with shRNAs targeting genes identified in the genome-wide CRISPR screen. Cells were treated with bafilomycin A1 to allow the accumulation of membrane-enclosed HaloTag7–LC3 and were sequentially labeled with the membrane-impermeant and membrane-permeant HaloTag ligands, as described for the Takahashi assay in Fig. 1C. The proportion of cells exhibiting increased MPL fluorescence relative to the peak of the shScramble control population was calculated. Candidates with an increased proportion of cells in this population were selected for secondary analysis in (B). (B) HeLa cells expressing HaloTag7–LC3 were transduced with the indicated shRNAs lentivirus, starved in EBSS for 3 or 24 h, pulse-labeled with TMR HaloTag ligand for 20 min, and chased in EBSS for 2 h. HaloTag7–LC3 degradation was analyzed by immunoblotting. Data represent three independent experiments, and the degradation ratio was calculated as indicated on the y axis. To minimize false-negative selection, shRNAs targeting the same candidate gene (nos. 7, 8, 16, 25, and 26) were retained for further validation, even when individual shRNAs, including nos. 7 and 25, did not show a statistically significant difference from the control. Statistical significance was assessed by one-way ANOVA (*p* < 0.05; \**p* < 0.01; \*\*\**p* < 0.0001). (C) Representative images of MPL fluorescence in cells expressing shRNAs nos. 16, 25, and 26. These cells exhibited fewer MPL- positive puncta than shScramble control cells. Scale bar, 10 μm.

**Figure S2.**
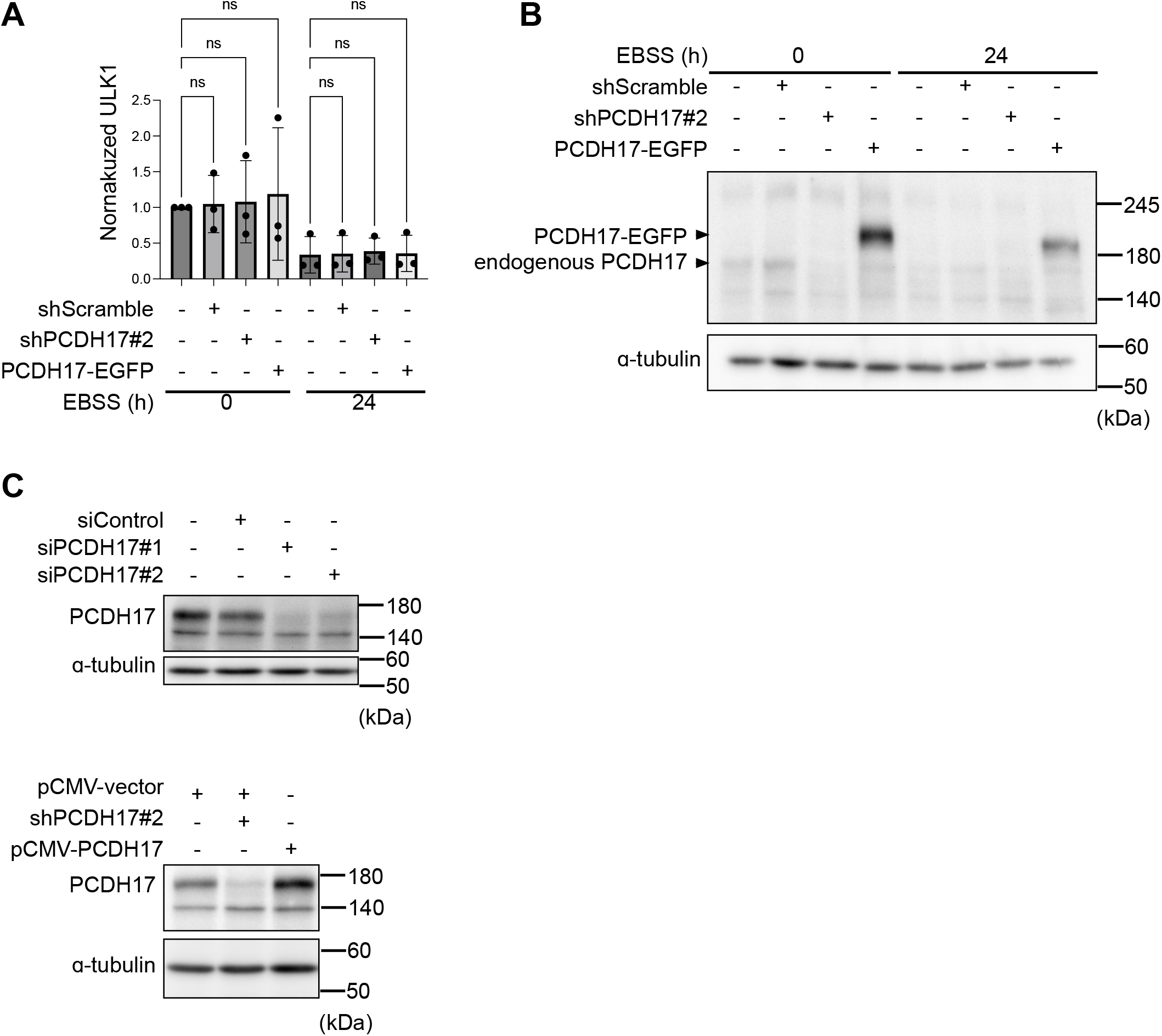
Validation of PCDH17 expression and analysis of ULK1 abundance. (A) ULK1 abundance in the immunoblots shown in Fig. 4A was quantified. Data are presented as the mean ± SD from three independent immunoblots. (B) Scramble control or PCDH17 knockdown or PCDH17–EGFP expressing Hela cells were maintained in complete medium or starved in EBSS for 24 h. Cell lysates were analyzed by immunoblotting with the anti-PCDH17 (C-region) or internal control antibodies. (C) Validation of PCDH17 depletion and re-expression. HeLa cells were transfected with siControl, siPCDH17#1, or siPCDH17#2 siRNA for 24 h. In parallel, parental HeLa cells or HeLa cells transduced with shPCDH17#2 lentivirus were transfected with pCMV vector or pCMV-PCDH17 plasmid, as indicated. Cell lysates were analyzed by immunoblotting with the anti-PCDH17 (C-region) or internal control antibodies.

**Figure S3.**
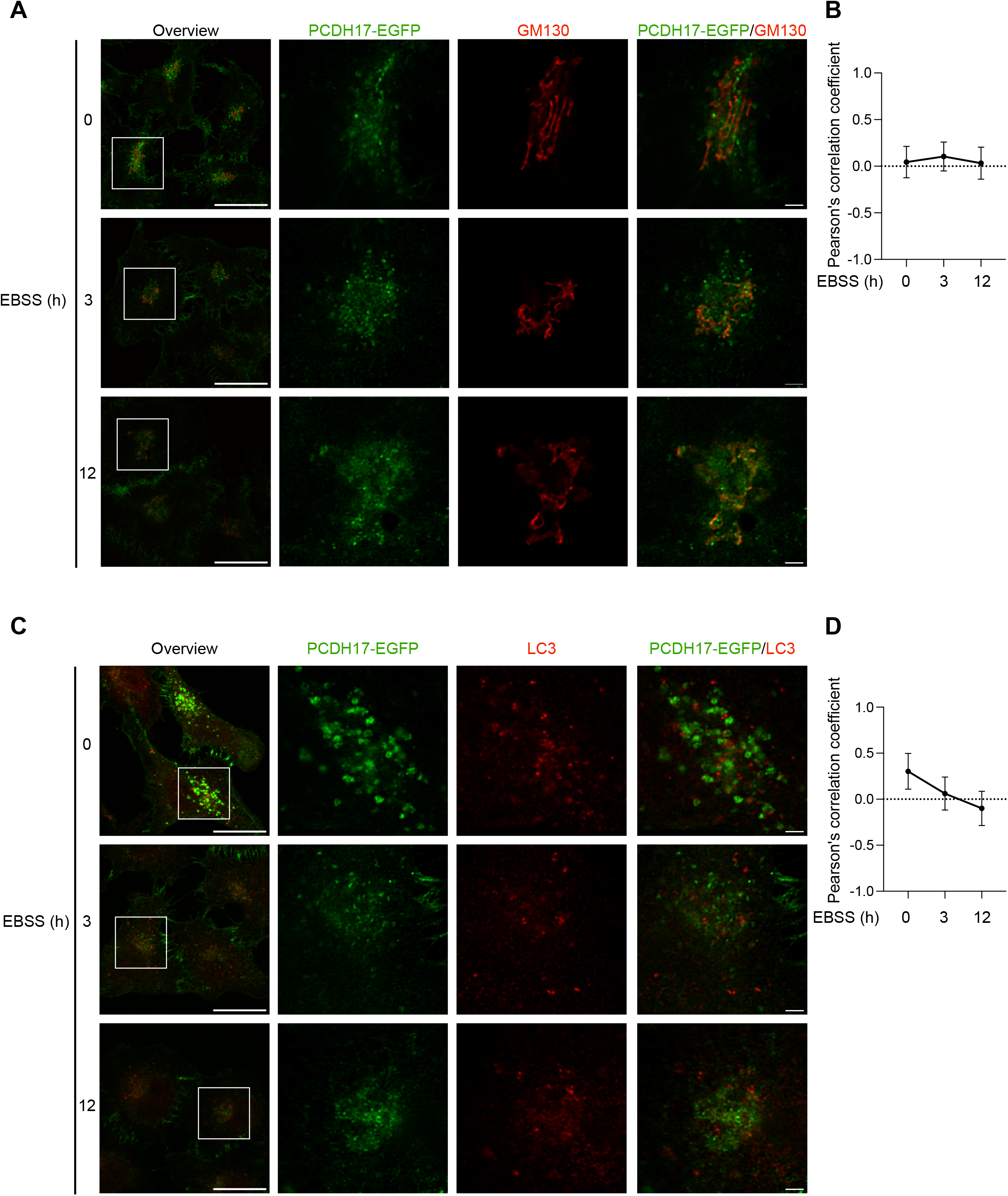
Starvation-dependent colocalization of PCDH17–EGFP with GM130 and LC3. (A, C) HeLa cells expressing PCDH17–EGFP were starved in EBSS for the indicated durations, fixed, and stained with antibodies against GM130 (A) or LC3 (C). Boxed regions in the overview images are enlarged in the adjacent panels. The enlarged images were deconvolved using Huygens software. Scale bars, 20 μm in the overview images and 2 μm in the enlarged images. (B, D) Colocalization of PCDH17–EGFP with GM130 (B) or LC3 (D) was quantified as Pearson’s correlation coefficients using the Coloc2 plugin in ImageJ. Pearson’s correlation coefficients calculated for pixels above the Costes threshold are shown (n > 20 cells).

**Figure S4.**
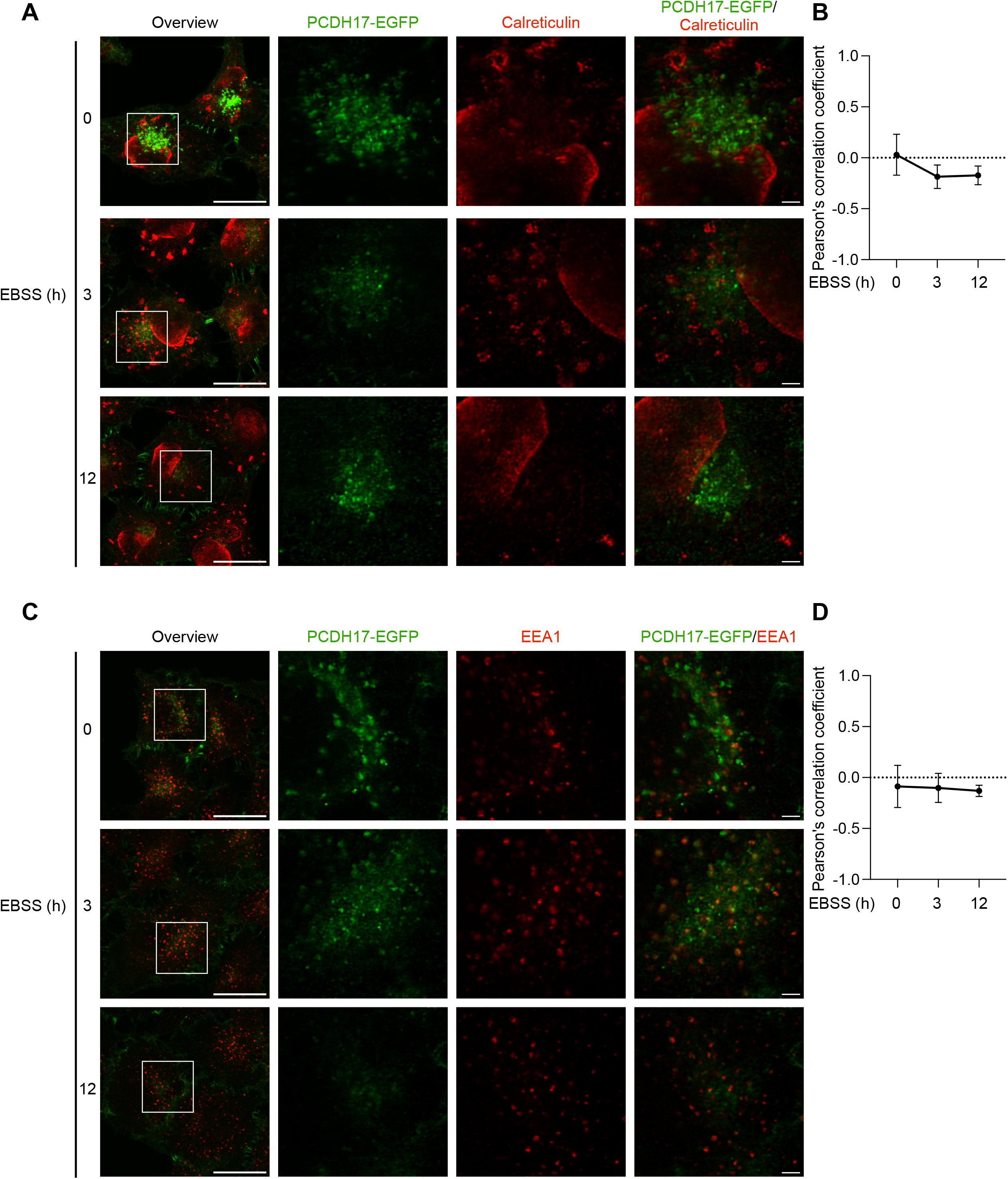
Starvation-dependent colocalization of PCDH17–EGFP with calreticulin and EEA1. (A, C) HeLa cells expressing PCDH17–EGFP were starved in EBSS for the indicated durations, fixed, and stained with antibodies against calreticulin (A) or EEA1 (C). Boxed regions in the overview images are enlarged in the adjacent panels. The enlarged images were deconvolved using Huygens software. Scale bars, 20 μm in the overview images and 2 μm in the enlarged images. (B, D) Colocalization of PCDH17–EGFP with calreticulin (B) or EEA1 (D) was quantified as Pearson’s correlation coefficients using the Coloc2 plugin in ImageJ. Pearson’s correlation coefficients calculated for pixels above the Costes threshold are shown (n > 20 cells).

**Figure S5.**
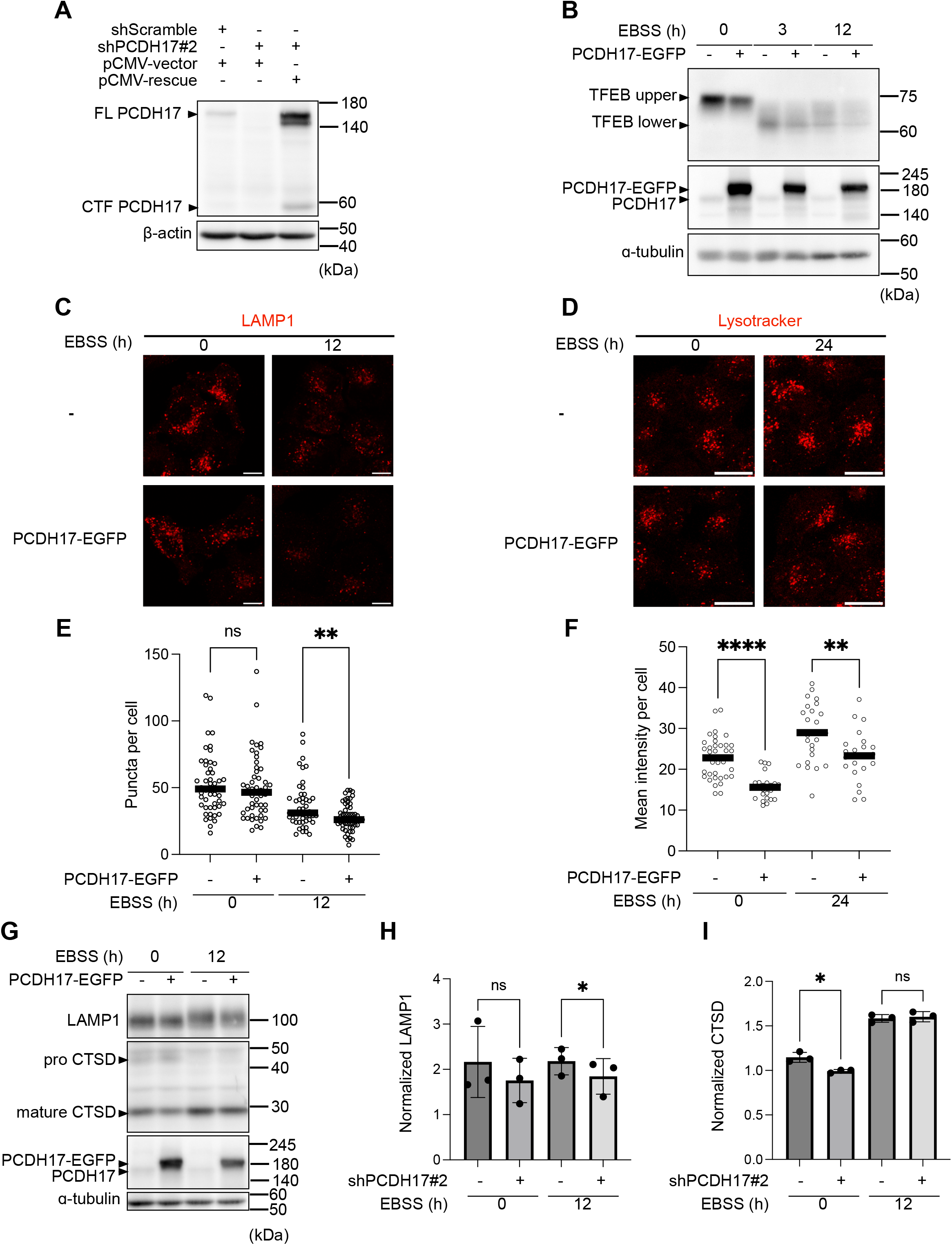
Expression of shRNA-resistant PCDH17 and lysosomal phenotypes induced by PCDH17 overexpression. (A) Scramble control or PCDH17 knockdown Hela cells were transfected with pCMV vector or shRNA-resistant PCDH17 plasmid (pCMV-rescue) for 24 h. PCDH17 expression was assessed by immunoblotting. (B) HeLa cells with or without PCDH17-EGFP expression were maintained in complete medium or starved in EBSS for 3 or 12 h. Cell lysates were analyzed by immunoblotting with antibodies against PCDH17 (C-region), TFEB and the indicated loading control. (C, D) HeLa cells with or without PCDH17-EGFP expression were maintained in complete medium or starved in EBSS for 12 h (C) or 24 h (D). Cells were fixed and processed for immunofluorescence analysis of LAMP1-positive puncta (C), or stained with LysoTracker before live-cell imaging (D). Scale bars, 10 μm in (C) and 20 μm in (D). (E, F) Quantification of the number of LAMP1-positive puncta per cell (E) and LysoTracker fluorescence (F) shown in (C) and (D), respectively. More than 45 cells were analyzed for LAMP1-positive puncta, and more than 20 cells were analyzed for LysoTracker fluorescence. (G–I) HeLa cells with or without PCDH17-EGFP expression were maintained in complete medium or starved in EBSS for 12 h. Cell lysates were analyzed by immunoblotting with the indicated antibodies (n = 3 independent experiments). Statistical significance was assessed by one-way ANOVA (\**p* < 0.05; \*\**p* < 0.01; \*\*\*\**p* < 0.0001).

**Table S1. Gene list analyzed in three independent genome-wide CRISPR screens.**

**Table S2. Gene symbols, catalog numbers, and target sequences of shRNAs used for secondary screening of candidate genes.**

**Video S1. Live-cell imaging of PCDH17–EGFP and LAMP1–mRFP under nutrient-rich conditions.**

HeLa cells stably coexpressing PCDH17–EGFP and LAMP1–mRFP were imaged by live-cell microscopy. Images were acquired at 2.58-s intervals for a total duration of 28.38 s. The yellow circle highlights a ring-like PCDH17–EGFP structure associated with, and moving synchronously with, an LAMP1–mRFP-positive compartment. Scale bar, 5 μm.

